# Diverse logics and grammar encode notochord enhancers

**DOI:** 10.1101/2022.07.25.501440

**Authors:** Benjamin P Song, Michelle F Ragsac, Krissie Tellez, Granton A Jindal, Jessica L Grudzien, Sophia H Le, Emma K Farley

## Abstract

The notochord is a key structure during chordate development. We have previously identified several enhancers regulated by Zic and ETS that encode notochord activity within the marine chordate *Ciona robusta (Ciona)*. To better understand the role of Zic and ETS within notochord enhancers, we tested 90 genomic elements containing Zic and ETS sites for expression in developing *Ciona* embryos using a whole-embryo, massively parallel reporter assay. We discovered that 39/90 of the elements were active in developing embryos; however only 10% were active within the notochord, indicating that more than just Zic and ETS sites are required for notochord expression. Further analysis revealed notochord enhancers were regulated by three groups of factors: (1) Zic and ETS, (2) Zic, ETS and Brachyury (Bra), and (3) Zic, ETS, Bra and FoxA. One of these notochord enhancers, regulated by Zic and ETS, is located upstream of *laminin alpha*, a gene critical for notochord development in both *Ciona* and vertebrates. Reversing the ETS sites in this enhancer greatly diminish expression, indicating that enhancer grammar is critical for enhancer activity. Strikingly, we find clusters of Zic and ETS binding sites within the introns of mouse and human *laminin alpha 1* with conserved enhancer grammar. Our analysis also identified two notochord enhancers regulated by Zic, ETS, FoxA and Bra binding sites: the Bra Shadow (BraS) enhancer located in close proximity to *Bra*, and an enhancer located near the gene *Lrig*. Randomizing the BraS enhancer demonstrates that although the Zic and ETS sites are necessary for enhancer activity, they are not sufficient. We find that FoxA and Bra sites contribute to BraS enhancer activity. Zic, ETS, FoxA and Bra binding sites occur within the *Ciona* Bra434 enhancer and vertebrate notochord *Brachyury* enhancers, suggesting a conserved regulatory logic. Collectively, this study deepens our understanding of how enhancers encode notochord expression, illustrates the importance of enhancer grammar, and hints at the conservation of enhancer logic and grammar across chordates.

## INTRODUCTION

Enhancers are genomic elements that act as switches to ensure the precise patterns of gene expression required for development (Levine, 2010). Enhancers regulate the timing, locations and levels of expression by binding of transcription factors (TFs) to sequences within the enhancer known as transcription factor binding sites (TFBSs) (Heinz et al., 2010; Liu and Posakony, 2012; Small et al., 1992; Spitz and Furlong, 2012; Swanson et al., 2010). This binding, along with protein-protein interactions, leads to recruitment of transcriptional machinery and activation of gene expression. While we understand that TFBSs regulate enhancers and mediate tissue-specific expression, we have limited understanding of how the sequence of an enhancer encodes a particular expression pattern and what combinations of binding sites within enhancers are able to mediate enhancer activity. Given that the majority of variants associated with disease and phenotypic diversity lie within enhancers (Maurano et al., 2012; Tak and Farnham, 2015; Visel et al., 2009), it is critical that we understand how the underlying enhancer sequence encodes tissue-specific expression and what types of changes within an enhancer sequence can cause changes in expression, cellular identity and phenotypes. A set of grammatical rules that define how enhancer sequence encodes tissue-specific expression is an attractive idea first suggested almost 30 years ago (Arnone and Davidson, 1997; Barolo, 2016; Levo and Segal, 2014; Thanos and Maniatis, 1995). The hypothesis for grammatical rules is based on the fact that proteins and the enhancer DNA have physical properties. These physical constraints govern the interaction of proteins with DNA and could be read out within the DNA sequence at the level of TFBSs. Enhancer grammar is composed of constraints on the number, type, and affinity of TFBSs within an enhancer and the relative syntax of these sites (orders, orientations, and spacings).

We previously identified grammatical rules governing notochord enhancers regulated by Zic and ETS TFBSs (Farley et al., 2016). We found that there was an interplay between affinity and organization of TFBSs, such that organization could compensate for poor affinity and vice versa. Using these rules, we identified two novel notochord enhancers, Mnx and Bra Shadow (BraS). These enhancers use low-affinity ETS sites in combination with Zic sites to encode notochord expression (Farley et al., 2016). Here, we focus on obtaining a deeper understanding of how enhancers regulated by Zic and ETS encode notochord expression.

Zic and ETS are co-expressed in the developing notochord (Figure 1). The notochord is a key feature of chordates and acts as a signaling center to pattern the neighboring neural tube, paraxial mesoderm, and gut (Herrmann and Kispert, 1994; Stemple, 2005). Specification of the notochord by Brachyury (Bra), also known as T, is highly conserved across chordates (Chesley, 1935; Chiba et al., 2009; Wilkinson et al., 1990; Yasuo and Satoh, 1993). Other TFs important for activation of notochord gene expression include Zic (Elms et al., 2004; Imai et al., 2002b; Kumano et al., 2006; Matsumoto et al., 2007; Yagi et al., 2004), ETS, which is a TF downstream of FGF signaling (Imai et al., 2002a; Matsumoto et al., 2007; Miya and Nishida, 2003; Schulte-Merker and Smith, 1995; Yasuo and Hudson, 2007), and FoxA (Ang and Rossant, 1994; Dal-Pra et al., 2011; José-Edwards et al., 2015; Katikala et al., 2013; Passamaneck et al., 2009; Weinstein et al., 1994).

**Figure 1.**
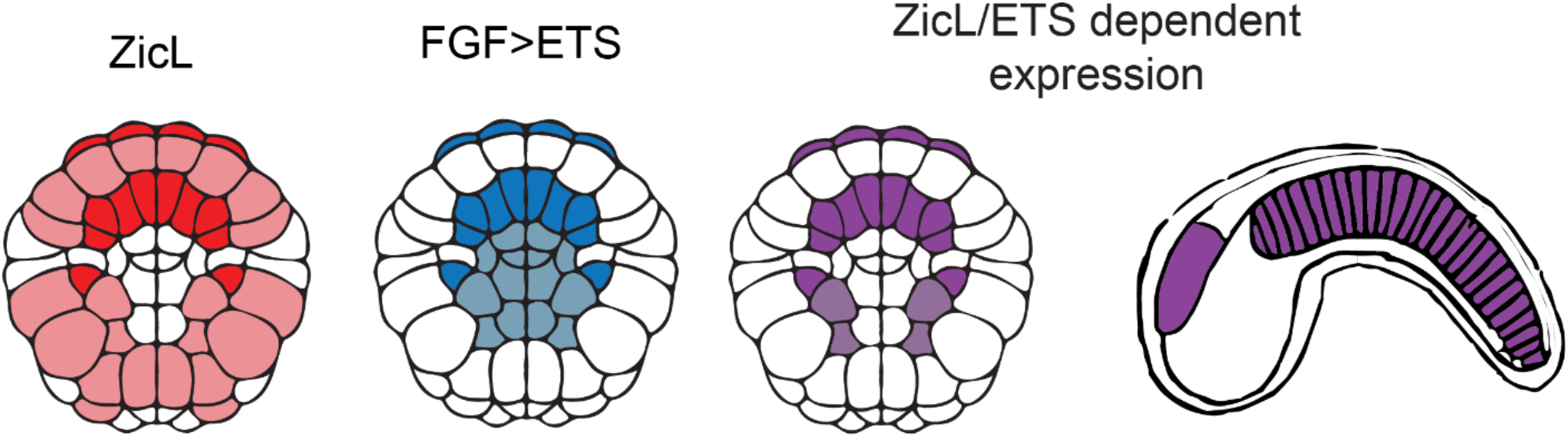
Zic and ETS expression in the 110-cell stage embryo. A. Co-expression of Zic and ETS is shown in purple and occurs in the notochord and a6.5 lineage, which gives rise to the anterior sensory vesicle and palps. A schematic of the tailbud embryos shows these cell types at a later stage of development. Dark coloring represents a6.5 and notochord lineages, light coloring represents other tissues with expression.

Our study focuses on the marine chordate, *Ciona*, a member of the urochordates, the sister group to vertebrates (Delsuc et al., 2006). Fertilized *Ciona* eggs can be electroporated with many enhancers in a single experiment which allows for testing of many enhancers in whole, developing embryos (Davidson and Christiaen, 2006). Furthermore, these embryos are easy to image, making it easy to determine the location of enhancer activity. These advantages, along with the fast development of *Ciona* and the similarity of notochord development programs between *Ciona* and vertebrates (Davidson and Christiaen, 2006), make it an ideal organism to study the rules governing notochord enhancers during development.

Within the *Ciona* genome, we found 1092 elements containing one Zic site and at least two ETS sites within 30bp of the Zic site. We tested 90 of these for expression in developing *Ciona* embryos. Only 10% of these regions drive notochord expression. These notochord enhancers fall into three categories: enhancers containing Zic and ETS sites, ones with Zic, ETS and Bra sites, and ones with Zic, ETS, FoxA and Bra sites. Within enhancers containing Zic and ETS sites, the organization of sites is important for activity, indicating that grammatical constraints on Zic and ETS encode enhancer activity. We find that one of the Zic and ETS enhancers is near an important notochord gene, *laminin alpha* (Veeman et al., 2008). The orientation of binding sites within this *laminin alpha* enhancer is critical for enhancer activity demonstrating the role of enhancer grammar. We find similar clusters of Zic and ETS sites within the introns of *laminin alpha-1* in both mouse and human. Strikingly, we find the same 12bp spacing between the Zic and ETS conserved across all three species. Additionally, this study identified two enhancers using a combination of Zic, ETS, FoxA, and Bra to encode notochord expression. One of these is the BraS enhancer. By creating a library of 25 million enhancer variants with only fixed Zic and ETS sites, we discover that although the Zic and ETS sites are necessary for enhancer activity, they are not sufficient,. We find that a Bra and FoxA site also contribute to activity of this enhancer. Other known *Bra* enhancers within Ciona (Corbo et al., 1997) and vertebrates (Schifferl et al., 2021) also harbor this combination of TFs, suggesting that Zic, ETS, FoxA, and Bra is a common feature of *Bra* regulation in chordates. Collectively, our study finds that grammar is a key component of functional enhancers with signatures of this enhancer logic and grammar seen across chordates, and provides a deeper understanding of *Bra* enhancers.

## RESULTS

### Searching for clusters of Zic and ETS sites within the *Ciona* genome

To better understand how Zic and ETS sites within enhancers encode notochord expression, we identified genomic regions containing one Zic site and at least two ETS sites within the 30 bp of the Zic site. As we have previously found that low-affinity ETS sites are required to encode notochord-specific expression (Farley et al., 2016), we searched for the core motif of ETS, GGAW (GGAA or GGAT), to consider all ETS sites regardless of affinity. We identified 1092 regions with a Zic and two ETS sites. To define Zic sites, we use EMSA and enhancer mutagenesis data from previous studies (Matsumoto et al., 2007; Takahashi et al., 1999; Yagi et al., 2004). The genomic regions were approximately 68bp in length. We define these regions as ZEE elements.

### Testing ZEE genomic elements for enhancer activity in developing *Ciona* embryos

We selected 90 ZEE elements (Figure S1 and Table S1) and synthesized these upstream of a minimal promoter (bpFog) and a transcribable barcode to conduct an enhancer screen (experiment outlined in Figure 2A). Each enhancer was associated with, on average, six unique barcodes. Each different barcode is a distinct measurement of enhancer activity. We electroporated this library into fertilized *Ciona* eggs. We collected embryos at the late gastrula stage when notochord cells are developing (Jiang and Smith, 2007) and both Zic and ETS are expressed (Imai et al., 2004; Winkley et al., 2021) (5.5 hours post fertilization, hpf). At this timepoint, we isolated mRNA and DNA. To determine that all the enhancer plasmids got into the embryos, we isolated the plasmids from the embryos and sequenced the DNA barcodes. We detected barcodes associated with all 90 ZEE elements from the isolated plasmids, indicating that all elements were tested for activity.

**Figure 2.**
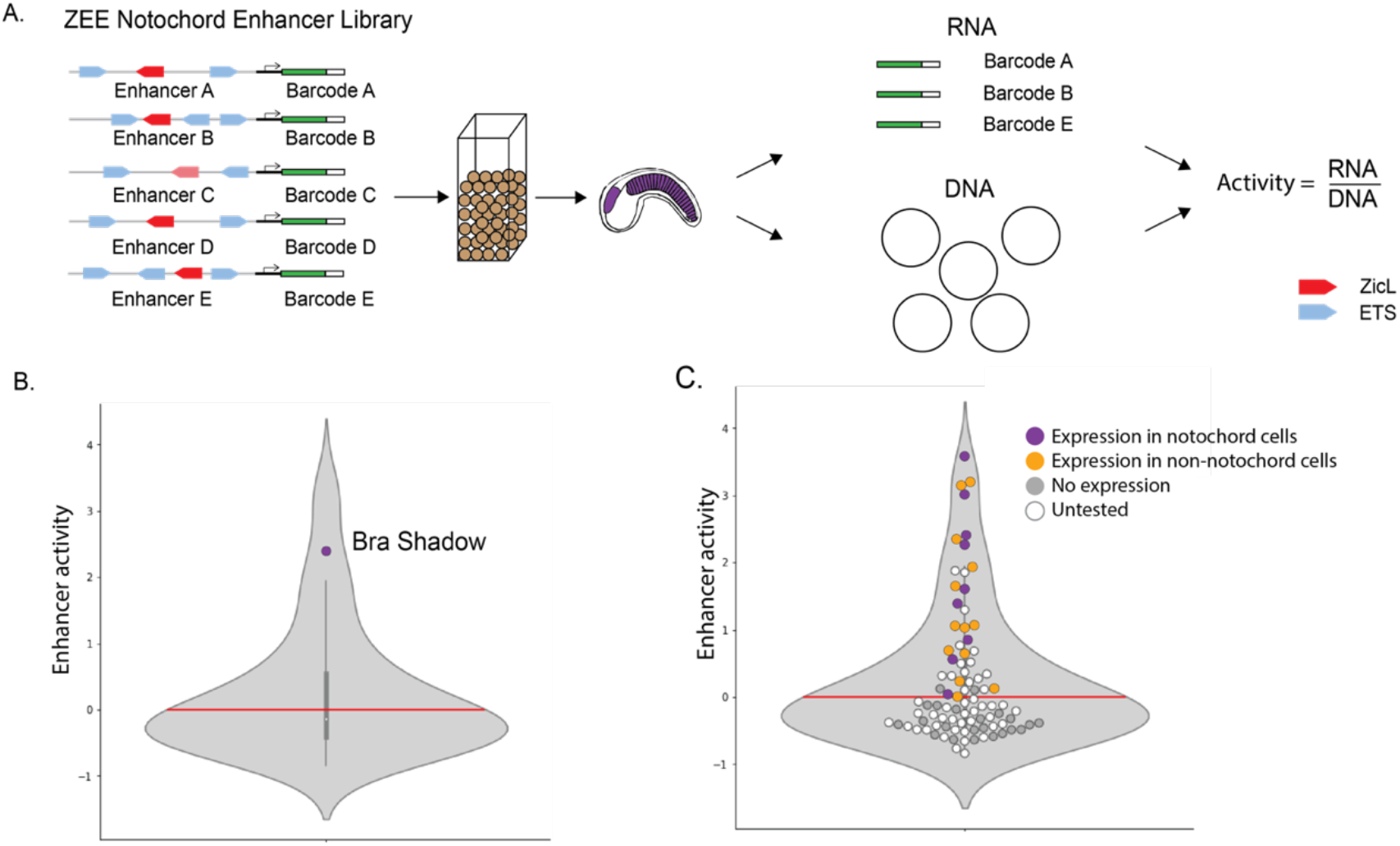
Screening Zic and ETS genomic elements in *Ciona*. A. Schematic of enhancer screen. 90 ZEE genomic regions, each associated with on average six unique barcodes were electroporated into fertilized *Ciona* eggs. mRNA and plasmid DNA were extracted from 5.5hpf embryos (tailbud embryo shown to highlight tissues with predicted expression). The mRNA barcodes and DNA barcodes were sequenced, and a normalized enhancer activity score was calculated for each enhancer by dividing the mRNA activity for a given enhancer by the number of copies of the plasmid. B. Violin plot showing the distribution of enhancer activity. The Bra Shadow enhancer served as a positive control and is labeled. The red line indicates the cut-off for non-functional elements at zero. C. Same plot as B, but with all 90 ZEE elements plotted as dots. Dots are colored by the results of an orthogonal screen, where we measured the GFP expression in at least 100 embryos to determine the location of expression. Enhancers driving notochord expression are shown in purple, enhancers with expression but no notochord expression are shown in orange. ZEE elements that do not drive expression are grey and untested enhancers are shown in white.

We next wanted to see how many of the 90 ZEE elements act as enhancers to drive transcription. Active enhancers will transcribe the GFP and the barcode into mRNA. To find the functional enhancers, we isolated the mRNA barcodes from our library and sequenced them. We analyzed the sequencing data and measured the reads per million (RPM) for each barcode. To calculate an average RPM for a given enhancer, we averaged the RPM for each barcode associated with an enhancer. To normalize the enhancer activity to the differences in the amount of plasmid and therefore number of copies of the enhancer electroporated into embryos, we took the log2 of the average enhancer activity divided by the DNA RPM for the same enhancer to create an enhancer activity score. Enhancer activity scores below zero are non-functional, while elements with scores above zero are considered functional enhancers. The highest activity score is around four. There was a high correlation between all three biological replicates (Figure S2).

### Many of these genomic ZEE elements are not enhancers

As an internal, positive control in our enhancer screen, we included the Bra Shadow (BraS) enhancer. This enhancer drives expression in the notochord and weak expression in the a6.5 lineage, both locations that express Zic and ETS (Farley et al., 2016). The BraS enhancer activity score is 2.4 (Figure 2B), indicating that our library screen is detecting functional enhancers. Thirty-nine of the ZEE elements act as enhancers in our screen, while fifty-one of the ZEE elements drove no expression. This suggests that genomic elements containing a single Zic site and at least two Ets sites are not sufficient to drive expression in the notochord. To further validate our sequencing data and to determine the tissue specific location of the functional enhancers, we selected 20 non-functional elements and 24 functional enhancers from our screen to test by an orthogonal approach. Each of these ZEE elements were cloned upstream of a minimal bpFog promoter and GFP. We electroporated each enhancer into fertilized eggs and analyzed the GFP expression of these ZEE elements at 8hpf in at least 150 embryos across three biological replicates. Collectively, we analyzed expression of these elements in over 6600 embryos with this orthogonal approach.

All 20 ZEE elements defined as non-functional in our library drove no GFP expression, validating our enhancer activity score cut off that we defined for non-functional enhancers (Fig 2C). As Zic and ETS are expressed in many cell types, including the muscle, endoderm, ectoderm, notochord and neural cell types (Hudson et al., 2016, 2007; Imai et al., 2006; Picco et al., 2007; Wagner and Levine, 2012), we expected to see that some functional enhancers may have expression in these cell types. However, we anticipated that enhancers under combinatorial control of ZEE sites may drive expression in the notochord and the a6.5 neural lineage, that gives rise to the neural cell types called the anterior sensory vesicle and the palps, where Zic and ETS are both expressed (Ikeda and Satou, 2016; Matsumoto et al., 2007; Wagner and Levine, 2012) (Figure 1). In the 24 enhancers detected as functional, 92% of these enhancers (22/24) showed GFP expression within the embryos. Nine drove expression in the notochord (Figure S3). Four of the enhancers are active almost exclusively in the notochord with some a6.5 expression (ZEE10, 13, 20, 27). The remaining five are active in the notochord and other cell types such as the endoderm. Twelve of the ZEE enhancers drove varying levels of expression in the a6.5 lineage, while one drove expression exclusively in this cell type (ZEE22). Thirteen drove expression in other cell types of the embryo (Table S2). These results indicate that our enhancer screen accurately detects functional enhancers, and our tissue-specific analysis provides detailed expression patterns for these enhancers.

### Elucidating the logic of the enhancers driving notochord expression

Having seen that so few enhancers drive expression in the notochord, we were interested to better understand why these nine functional enhancers were active in the notochord. It is possible that they are functional due to the grammar of the Zic and ETS sites or because other TFBSs are required for notochord expression. To investigate these two hypotheses, we looked at the nine notochord enhancers in more detail. FoxA and Bra are two other TFs important for activation of notochord enhancers (José-Edwards et al., 2015; Katikala et al., 2013; Passamaneck et al., 2009). We therefore searched all 90 ZEE elements for FoxA and Bra sites. We used TRTTTAY as the FoxA motif (Katikala et al., 2013; Li et al., 2017; Passamaneck et al., 2009) and TNNCAC as the Bra motif (Casey et al., 1998; Conlon et al., 2001; Di Gregorio and Levine, 1999; Dunn and Di Gregorio, 2009; Müller and Herrmann, 1997) based on EMSA and crystal structures.

### The nine elements that drive notochord expression contain three different combinations of transcription factors

Of the 90 genomic regions we tested, 42 had only Zic and ETS sites, 39 had Zic, ETS and Bra sites, 4 had Zic, ETS, FoxA, and Bra sites and 5 had Zic, ETS and FoxA sites. Ten percent of the enhancers containing only Zic and ETS sites drive notochord expression (4/42). Eight percent (3/39) of the enhancers containing Zic, ETS, and Bra drive notochord expression. None of the enhancers (0/5) containing Zic, ETS, and FoxA drive notochord expression, while fifty percent (2/4) of the enhancers containing Zic, ETS, FoxA and Bra are active in the notochord (Figure 3). Thus, there are three groups of notochord enhancers that contain: (1) Zic and ETS sites alone, (2) Zic, ETS and Bra sites, or (3) Zic, ETS, FoxA, and Bra sites. Having found that only a few of the elements containing Zic and ETS sites alone were functional, we wanted to understand if the organization or grammar of sites within these enhancers was important.

**Figure 3.**
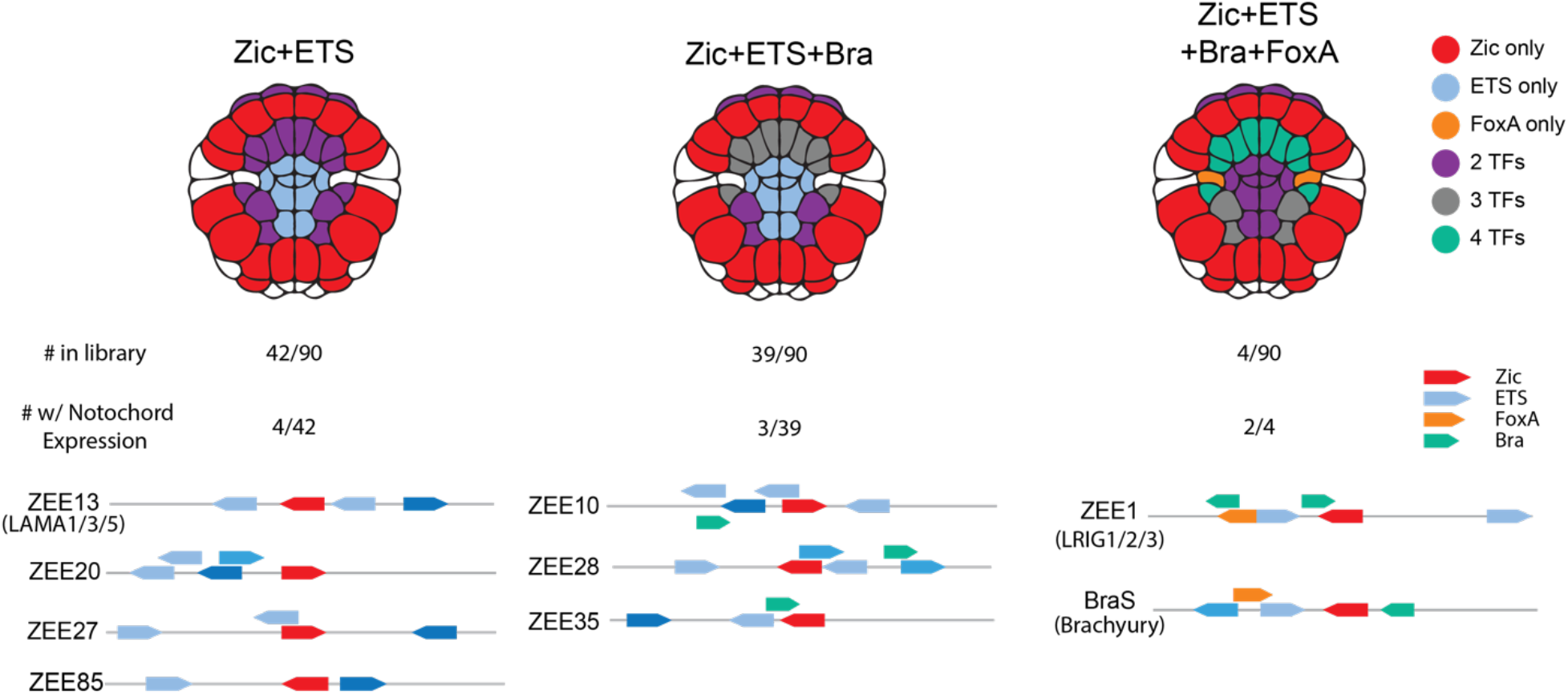
Combinations of transcription factors in ZEE enhancers that drive notochord expression. Notochord-expressing ZEE elements were grouped by the combination of transcription factor binding sites present in each element. For each combination, an embryo schematic shows the overlapping region of expression for that given combination. Below the embryo schematic, the number of ZEE elements, the number with notochord expression and a schematic of the ZEE elements with notochord expression for that combination of transcription factors. Zic (red), ETS (blue), FoxA (orange), and Bra (green) sites are annotated. Color saturation of ETS sites indicates binding site affinity (increasing affinity is indicated by an increase in color saturation).

### Zic and ETS enhancer grammar encodes notochord *laminin alpha* expression

Four ZEE enhancers driving notochord-specific expression contain Zic and ETS binding sites. Three of these elements are not in close proximity to known notochord genes, though it is possible that these elements regulate notochord genes further away. The ZEE13 enhancer is located close to *laminin alpha*, which is critical for notochord development (Veeman et al., 2008) (Figure 4A). Given the proximity of this notochord-specific enhancer to *laminin alpha*, we decided to focus further analysis on this enhancer, which we renamed the Lama enhancer. Notably this enhancer contains three ETS sites. To determine the affinity of these sites, we used Protein Binding Microarray data (PBM) for mouse ETS-1 (Wei et al., 2010), as the binding specificity of ETS is highly conserved across bilaterians (Nitta et al., 2015; Wei et al., 2010). The consensus highest-affinity site has a score of 1.0, and all other 8-mer sequences have a score relative to the consensus. The Lama enhancer contains two ETS sites with exceptionally low affinities of 0.10, or 10% of the maximal binding affinity, while the most distal ETS site is a high-affinity site (0.73).

**Figure 4.**
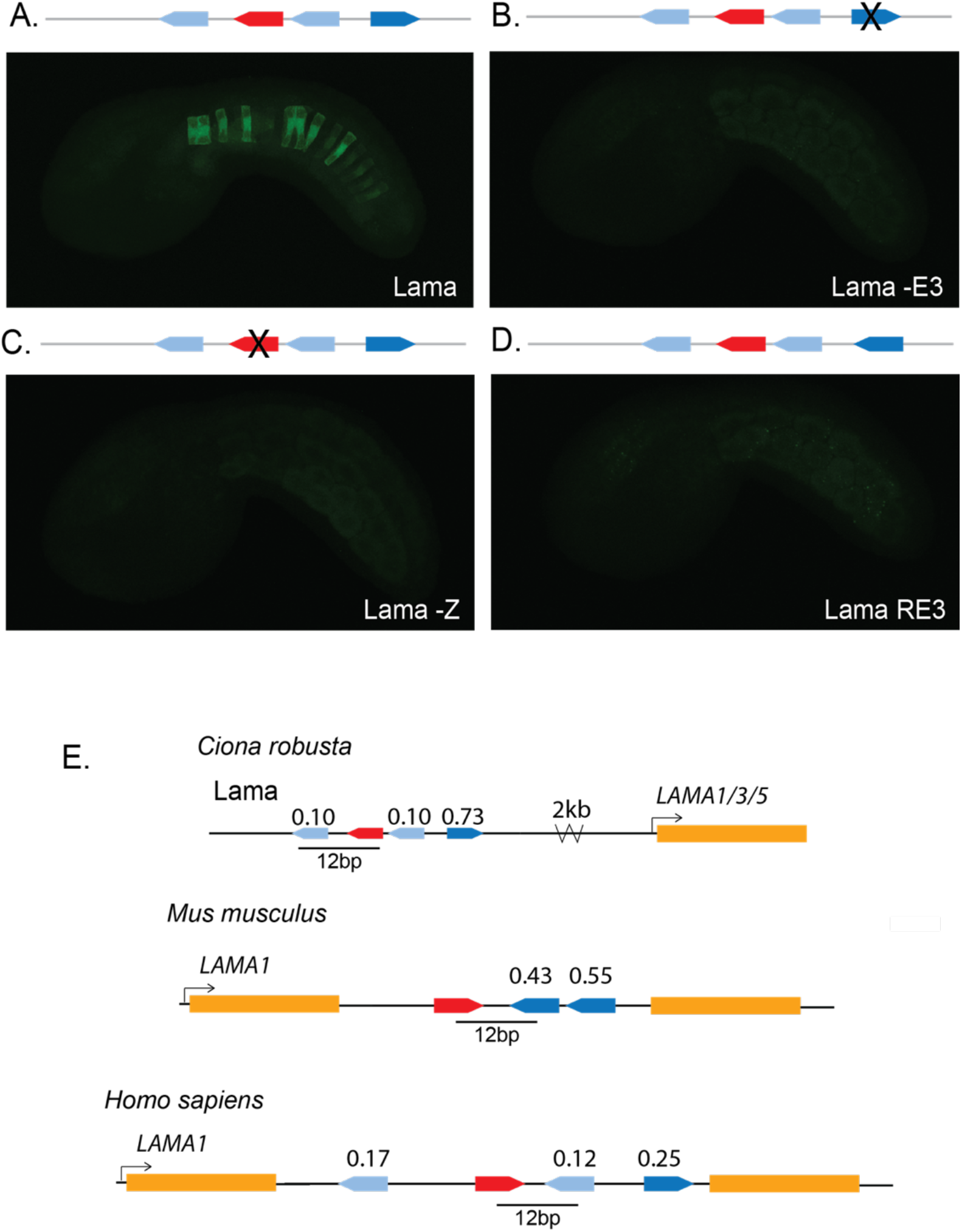
Zic and ETS grammar encodes a notochord *laminin alpha* enhancer. A. Embryo electroporated with the Lama enhancer (ZEE13); GFP expression can be seen in the notochord. B. Embryo electroporated with Lama -ETS3, where ETS3 was mutated to be non-functional; no GFP expression detected. C. Embryo electroporated with Lama -Z, where the Zic was mutated to be non-functional; no GFP expression detected. D. Embryo electroporated with Lama ETS3, where the sequence of ETS3 was mutated to be the reverse complement; no GFP expression detected. Comparable results were seen when ETS1 was reversed E. Schematics of Zic and ETS clusters in the genome of *Ciona*, mouse, and human. All three *laminin alpha-1* clusters have a spacing of 12bp between an ETS and Zic site and all contain non-consensus ETS sites. ETS site affinity scores are noted above each site. Color saturation of ETS sites indicates binding site affinity (increasing affinity is indicated by an increase in color saturation).

To determine if the Zic site and three ETS sites are important for enhancer activity, we made a point mutation to ablate the ETS3 site (Figure 4B, Figure S4A, and Table S3). This led to a complete loss of notochord activity. Similarly, ablation of the Zic site results in complete loss of enhancer activity, indicating that both Zic and ETS sites are necessary for activity of this Lama enhancer (Figure 4C). Previously, we saw that the organization of sites within enhancers, a component of enhancer grammar, is critical for enhancer activity in both the Mnx and Bra enhancer. To see if enhancer grammar is important for activity within the Lama enhancer, we altered the orientation of sites within this enhancer and measured the impact on enhancer activity. Reversing the orientation of the first ETS site, which has an affinity of 0.10, led to a dramatic reduction in notochord expression, suggesting the orientation of this ETS site is important for enhancer activity. Similarly, reversing the orientation of the third ETS site (Lama RE3), which has an affinity of 0.73, also causes a loss of notochord expression (Figure 4D, Figure S4A, and Table S3). These two manipulations demonstrate that the orientation of the ETS sites within this enhancer is important for activity, and thus, that there are some grammatical constraints on the *Ciona* Lama enhancer. It is likely that grammar is an important feature of enhancers regulated by Zic and ETS, as we have previously seen similar grammatical constraints on the orientation and spacing of binding sites within the Mnx and BraS enhancer.

### Vertebrate *laminin alpha* introns contain clusters of Zic and ETS with conserved spacing

The expression of laminin in the notochord is highly conserved between urochordates and vertebrates (Reeves et al., 2017; Scott and Stemple, 2005; Veeman et al., 2008). Indeed, laminins play a vital role in both urochordate and vertebrate notochord development, with mutations in laminins or components that interact with laminins causing notochord defects (Machingo et al., 2006; Parsons et al., 2002; Pollard et al., 2006). The *Ciona laminin alpha* is the ortholog of the vertebrate *laminin alpha 1/3/5* family.

We therefore sought to determine if we could find a similar combination of Zic and ETS sites in proximity to vertebrate *laminin* genes. Strikingly, we find a cluster of Zic and ETS sites within the intron of both the mouse and human *laminin alpha-1* gene. The affinity of the ETS sites in all three species is also far from the consensus: the human cluster contains three ETS sites of 0.12, 0.17 and 0.25 affinity, while the putative mouse enhancer contains fewer, but higher-affinity, ETS sites. We have previously seen that the spacing between Zic and adjacent ETS sites affects levels of expression, with spacings of 11 and 13bp seen between ETS and Zic sites in the BraS enhancer and Mnx enhancer, respectively (Farley et al., 2016). In line with this observation, the *laminin alpha-1* clusters in mouse and human and the *Ciona* Lama enhancer have a 12bp spacing between the ETS and adjacent Zic site in all three species, suggesting that such spacings (11-13bp) are a feature of some notochord enhancers regulated by Zic and ETS. The conservation of this combination of sites, the low-affinity ETS sites, and the conserved spacing hints at the conservation of enhancer grammar across chordates.

### The Zic, ETS, FoxA and Bra regulatory logic encodes notochord enhancer activity

The group of genomic elements most enriched in notochord expression was the group containing Zic, ETS, FoxA and Bra binding sites, with two of the four driving notochord expression. Both of these enhancers are located near genes expressed in the notochord (Reeves et al., 2017). The first was our positive control BraS, while the second enhancer is in proximity of the *Lrig* gene.

We previously identified the BraS enhancer through a search for rules governing Zic and ETS grammar that included number and type of TFBSs, along with the affinity, spacing, and orientation of TFBSs (Farley et al., 2016). The BraS enhancer contains a Zic and two low-affinity ETS sites (0.14 and 0.25). Changing the orientation of the lowest affinity ETS site, located 11bp from the Zic site, leads to loss of expression, indicating that there are grammatical constraints on this enhancer. To confirm the role of the Zic and two ETS sites within BraS, we ablated these three sites (Zic and both ETS sites) with point mutations; this leads to complete loss of expression, demonstrating that these sites are necessary for notochord expression (Figure 5B, Figure S4B, and Table S3). To test if these sites are sufficient for notochord expression, we randomized every nucleotide within the enhancer except for these three sites, creating an enhancer harboring a Zic and two ETS sites that are constant in sequence and position in a sea of 24.5 million enhancer variants. We electroporated this library into embryos and counted GFP expression in 8hpf embryos. The ZEE-randomized BraS enhancer drives significantly lower expression within the notochord than the BraS enhancer, indicating that there are other sites within the enhancer that are also important for tissue-specific expression (Figure 5C, Figure S4B, and Table S3). This experiment highlights the importance of understanding sufficiency in addition to necessity of sites.

**Fig 5.**
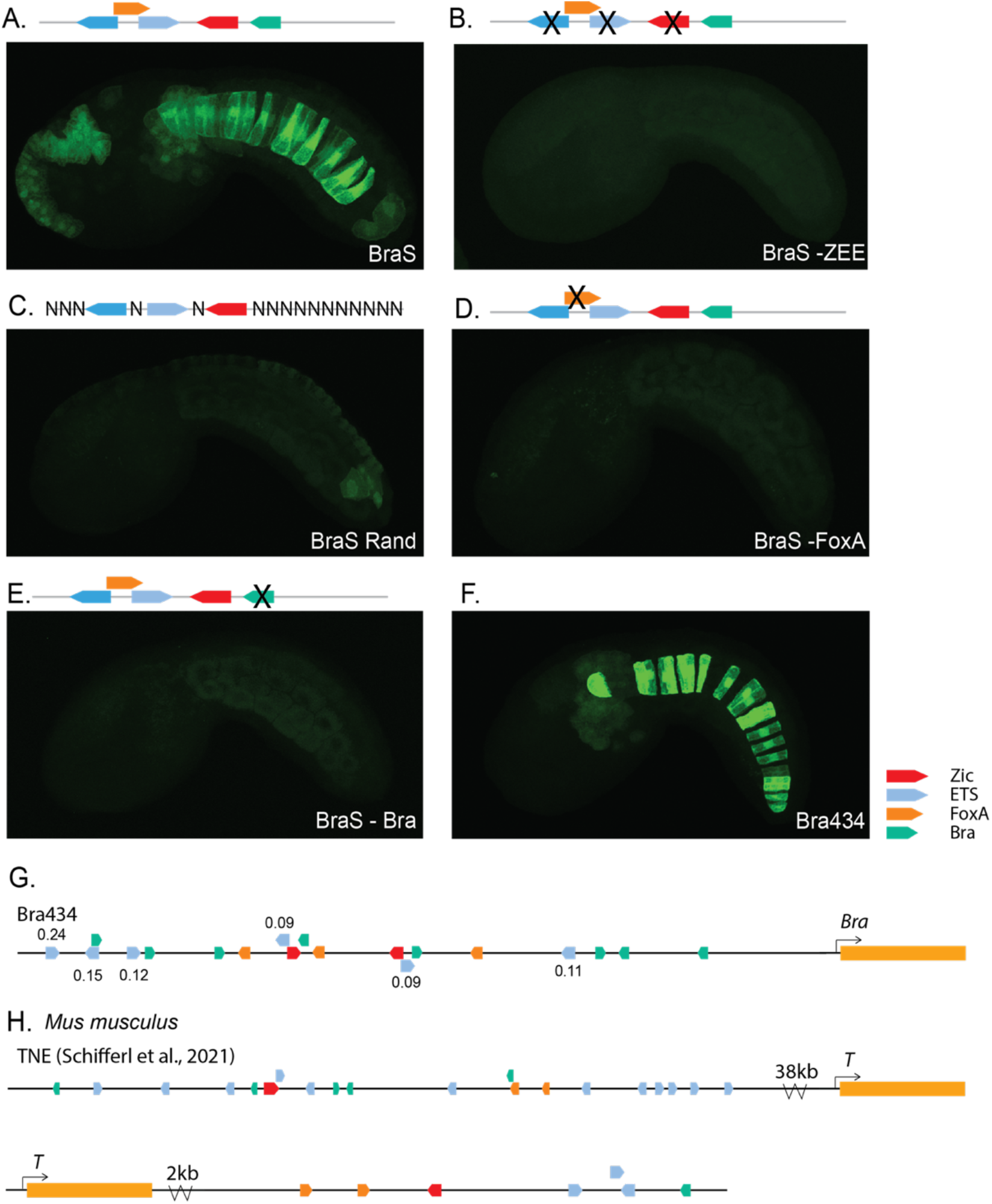
Zic, ETS, FoxA, and Bra may be a common regulatory logic for Brachyury enhancers. **A**. Embryo electroporated with the Brachyury shadow (BraS) enhancer; GFP expression can be seen in the notochord. **B**. Embryo electroporated with BraS -ZEE, where the Zic and two ETS sites were mutated to be non-functional; no GFP expression was detected. **C**. Embryo electroporated with BraS Rand, where the Zic and two ETS sites were fixed, and all other nucleotides were randomized; GFP expression was greatly diminished. **D**. Embryo electroporated with BraS -FoxA, where the sequence of FoxA was mutated to be non-functional; GFP expression was greatly diminished. **E**. Embryo electroporated with BraS -Bra, where the sequence of Bra was mutated to be non-functional; GFP expression was greatly diminished. **F**. Embryo electroporated with Bra434; GFP expression can be seen in the notochord. G-I. Schematics of Zic (red), ETS (blue), Fox (orange), and Bra (green) clusters near *Bra* in the genomes of *Ciona* and mouse.

Two obvious candidates for additional functional sites within BraS are the FoxA and Bra sites, which we detected in this enhancer. Both FoxA and Bra are TFs known to regulate notochord enhancers in urochordates and vertebrates (Ikeda and Satou, 2016; José-Edwards et al., 2015; Kumano et al., 2006; Lolas et al., 2014; Passamaneck et al., 2009; Reeves et al., 2021). Ablating the Bra site within BraS leads to a reduction in expression, as does ablating the FoxA site (Figure 5D and E, Figure S4B, and Table S3). These manipulations suggest that all five sites (Zic, FoxA, Bra, and two ETS sites) are important for enhancer activity, and that all four TFs contribute to the activity of BraS. Studies of enhancers often stop when mutation experiments demonstrate a TF is required for enhancer activity. However, this falls short of a full understanding of these enhancers. Our results highlight that finding necessary sites is not enough to identify all functional sites within the enhancer. Indeed, we now have a deeper understanding of the BraS enhancer, namely that it is regulated by Zic, ETS, Bra and FoxA.

### Zic, ETS, Bra and FoxA may be a common regulatory logic for *Ciona Brachyury* enhancers

The first and most well-studied *Bra* enhancer is the Bra434 enhancer (Corbo et al., 1997; Fujiwara et al., 1998), which drives strong expression in the notochord (Figure 5F). The Bra434 enhancer contains Zic, ETS, FoxA, and Bra sites; ablating these sites within this enhancer lead to reduced expression, suggesting that these sites contribute to enhancer activity. There are different reports regarding the number and location of Zic, ETS, FoxA, and Bra sites within the Bra434 enhancer depending on the method used to define sites (Corbo et al., 1997; Shimai and Veeman, 2021). Here we annotate the Bra434 enhancer using crystal structure data, enhancer mutagenesis data, EMSA and PBM data. Our approach identifies two Zic sites, six ETS sites, three FoxA sites, and eight Bra sites (Figure S5). Of these TFs, the least information is available regarding Zic; thus it is possible that there are other more degenerate Zic sites that may be identified in future studies. (Corbo et al., 1997; Fujiwara et al., 1998; Reeves et al., 2021; Shimai and Veeman, 2021). Having seen that clusters of Zic, ETS, FoxA, and Bra are important in the BraS and Bra434 enhancers, we next wanted to see if this logic is found in *Bra* enhancers in vertebrates.

### Vertebrate notochord enhancers contain clusters of Zic, ETS, Fox and Bra, suggesting this is a common mechanism for regulation of *Brachyury* expression in the notochord

In mouse, the most well-defined notochord enhancer to date is within an intron of *T2*, 38kb upstream of *T*, which is the mouse ortholog of *Bra* (Schifferl et al., 2021). This mouse *T* enhancer is required for *Bra*/*T* expression, notochord cell specification and differentiation. Homozygous deletion of this *Bra*/*T* enhancer in mouse leads to reduction of *Bra*/*T* expression, a reduction in the number of notochord cells, and halving of tail length. Bra/T and FoxA binding sites have previously been identified within this enhancer. We find that this mouse *Bra*/*T* enhancer also contains Zic and ETS binding sites. Within this enhancer there are 12 ETS sites; 11 of these have affinities ranging from 0.09-0.14, while one site has an affinity of 0.65, indicating that this enhancer contains low-affinity ETS sites.

As we saw with the *Ciona* BraS and Bra434 enhancer, typically there are multiple enhancers that all regulate the same or similar patterns of expression (Frankel et al., 2010; Hong et al., 2008; Perry et al., 2010). This is thought to confer the transcriptional robustness required for successful development (Antosova et al., 2016; Frankel et al., 2010; Osterwalder et al., 2018; Perry et al., 2010). Following this logic, we continued to search the mouse *T* region to see if we could find other putative notochord enhancers that may regulate *Bra*. We identified a region located 2kb downstream of *T* that contains a cluster of Zic, ETS, FoxA and Bra sites. This putative enhancer occurs within an open chromatin region in mouse E8.25 notochordal cells (Pijuan-Sala et al., 2020), suggesting this may be another mouse *T* enhancer. This second putative *T* enhancer contains three low-affinity ETS sites of 0.11, 0.11 and 0.12. Similarly in zebrafish, a notochord enhancer located 2.1kb upstream of the *Bra* ortholog *ntl* (Harvey et al., 2010) also contains a cluster of Zic, ETS, FoxA, and Bra sites. The presence of these four TFs in *Ciona*, zebrafish, and mouse *Bra* enhancers suggest that the use of Zic, ETS, FoxA and Bra could be a common enhancer logic regulating expression of the key notochord specification gene *Bra* in chordates.

## Discussion

In this study we sought to understand the regulatory logic of notochord enhancers by taking advantage of high-throughput studies within the marine chordate *Ciona*. Within the *Ciona* genome, there are 1092 genomic regions containing a Zic site within 30bp of two ETS sites. We tested 90 of these ZEE genomic regions for expression in developing *Ciona* embryos. Surprisingly, only nine of the regions drove notochord expression. Among these nine, we identified a *laminin alpha* enhancer that was highly dependent on grammatical constraints for proper expression. We found a similar cluster of Zic and ETS sites within the intron of the mouse and human *laminin alpha-1* gene, strikingly, these clusters and the *Ciona* laminin enhancer have the same spacing between the sites Zic and ETS sites. Within the Bra Shadow enhancer, although Zic and ETS are necessary for enhancer activity, randomization of BraS by keeping only the Zic and ETS sites constant in a sea of 24 million variants finds that these sites are not sufficient for notochord activity. FoxA and Bra sites are also necessary for notochord expression. This combination of sites occurs within other *Bra* enhancers in *Ciona* and vertebrates suggesting this combination of TFs may be a common logic regulating *Bra* expression. Our study identifies new developmental enhancers, demonstrates the importance of enhancer grammar within developmental enhancers and provides a deeper understanding of the regulatory logic governing *Brachyury*. Our findings of the same clusters of sites within vertebrates hint at the conserved role of grammar and logic across chordates.

### Very few genomic regions containing Zic and two ETS sites are functional enhancers

Our analysis of 90 genomic elements all containing one Zic site in combination with two ETS sites strikingly demonstrated that clusters of sites are not sufficient to drive expression. Only 39 of the 90 elements tested drove any expression, and even more surprisingly, only 15 of these drove expression in the predicted lineages, ASV or notochord. These findings indicate that searching for clusters of TFs is only minimally effective in identification of enhancers.

### Grammar is a key constraint of the Lama and BraS enhancers

The Lama enhancer is a ZEE element. Within the Lama enhancer, the orientation of binding sites relative to each other was critical for expression, providing evidence that enhancer grammar is a critical feature of functional enhancers regulated by Zic and ETS. Flipping the orientation of either the first or last ETS sites relative to the Zic site led to loss of enhancer activity in the *Ciona* Lama enhancer. This mirrors the results of flipping the orientation of the ETS sites within the BraS enhancer (Farley et al., 2016). *Laminin alpha* is a key gene involved in notochord development in both *Ciona* and vertebrates (Pollard et al., 2006; Veeman et al., 2008). Intriguingly, we find that both the human and mouse *laminin alpha-1* have introns that harbor a similar cluster of Zic and ETS sites to those seen within *Ciona*. There is a conservation of 12bp spacing between the Zic and ETS site, similar to the spacing we have observed between Zic and ETS sites within the notochord enhancers Mnx and BraS (Farley et al., 2016).

### Necessity of sites does not mean sufficiency – a deeper understanding of the BraS enhancer

Our study of the BraS enhancer highlights the importance of testing sufficiency of sites to investigate if we fully understand the regulatory logic of an enhancer. We previously demonstrated that reversing the orientation of an ETS site led to loss of notochord expression in the BraS enhancer. Here, in this study, we show via point mutations that both Zic and ETS sites are required for enhancer activity. However, randomization of the BraS enhancer to create 25 million variants in which the only the Zic and ETS sites are constant finds that these sites are not sufficient for enhancer activity, as the randomized BraS enhancer has reduced levels of enhancer activity. These sufficiency experiments are rarely done, and we are unaware of another study that has done this across the entirety of an enhancer. However, this experiment illustrates the importance of testing sufficiency to determine all the features contributing to enhancer function. Having discovered that Zic and ETS alone were not sufficient, we find that both FoxA and Bra sites also contribute to the enhancer activity, providing a deeper understanding of the regulation of the BraS enhancer.

### Limited binding site dependency information can provide signatures that identify enhancers, but improved understanding could lead to more accurate predictions

We were able to find the BraS enhancer using grammatical constraints on organization and spacing between Zic and ETS site and affinity of ETS sites. Interestingly, we did not have all the features required for enhancer activity. As such, this suggests that partial knowledge of grammatical constraints, or partial signatures of grammar could be used as to identify functional enhancers. Our previous strategy searched for these grammatical constraints in proximity of known notochord genes, which may be why we were successful in identification of the Mnx and BraS enhancer. However, the fact that we had not found all the features necessary and sufficient for enhancer activity could also explain why our search for elements containing ZEE had limited success. Understanding the dependency between all features within an enhancer will likely enable greater success in identification of functional regulatory elements. However, until then, our current knowledge of grammatical constraints may still be useful as a guide in pointing us toward putative enhancers

### Zic, ETS, FoxA, and Bra may be a common logic upstream of *Brachyury* in chordates

The Bra434 enhancer also contains the same combination of sites as the BraS enhancer; therefore, it is possible that this is a common logic for regulating *Bra*. Interestingly, we find these sites within mouse and zebrafish Brachyury enhancers (Harvey et al., 2010; Schifferl et al., 2021). While there are differences in expression dynamics of these factors in vertebrates and ascidians, it is striking to see this combination of sites in a validated notochord enhancers across these species. Indeed, our study in both the *laminin* enhancers and *Bra* enhancers provides hints of a conserved regulatory logic seen across chordates.

### Approaches to understanding dependency grammar of notochord expression

Searching for grammatical rules governing enhancers requires comparison of functional enhancers with the same features. Although we thought we had the same features in all 90 regions, we actually had at least three distinct types of enhancers within our screen. This illustrates a common problem in mining genomic data for patterns, as the assumption that we are comparing like with like is often an incorrect one. To uncover the grammatical constraints on enhancers, we need to not only understand the number and types of sites within an enhancer, but also the dependency between these sites, such as affinity, spacing, and orientation. Further screens with increased size and complexity and that combine both synthetic enhancers and genomic elements will likely be required to pinpoint the rules governing enhancer activity within genomes. Despite the complexity of studying enhancers in developing embryos, our study demonstrates that enhancer grammar is critical for encoding notochord activity and our observation of these logics and grammar signatures in vertebrates hints at conservation of these grammatical constraints across chordates.

## Author contributions

E.K.F., B.P.S., M.F.R, K.T., G.A.J. designed experiments. B.P.S., K.T., J.L.G., S.H.L. conducted experiments. M.F.R. conducted bioinformatic analyses. E.K.F. and B.P.S wrote the manuscript. All authors were involved in editing the manuscript.

## Acknowledgements

We thank the Farley lab for helpful discussions. We thank Janet H.T. Song for her critical reading of the manuscript. We thank the UCSD IGM Genomics Center for their assistance with sequencing.

## KEY RESOURCES TABLE

| REAGENT or RESOURCE | SOURCE | IDENTIFIER |
| --- | --- | --- |
| Deposited Data |  |  |
| snATACseq mouse E8.25 | Pijuan-Sala et al., 2020 | GEO: [GSE133244](https://www.ncbi.nlm.nih.gov/geo/query/acc.cgi?acc=GSE133244) |
| FACS-sorted notochord RNA-Seq | Reeves et al., 2017 | N/A |
| Human reference genome NCBI build 38, GRCh38 | Genome Reference Consortium | https://www.ncbi.nlm.nih.gov/grc/human |
| Mouse reference genome NCBI build 39, | Genome Reference Consortium | https://www.ncbi.nlm.nih.gov/grc/mouse |
| Ciona robusta genome | Satoh et al., 2019 | N/A |
| mouse ETS-1 universal PBM data | Wei et al., 2010 | https://thebrain.bwh.harvard.edu/uniprobe/index.php |
| ZEE library screen | This paper | N/A |
| Experimental Models: Organisms/Strains |  |  |
| Ciona intestinalis type A (Ciona robusta) | M-Rep | N/A |
| Oligonucleotides |  |  |
| Oligonucleotides for library screen, see Table S1 | This paper | N/A |
| Oligonucleotides for mutagenesis, see Table S4 | This paper | N/A |
| Recombinant DNA |  |  |
| Plasmid: BraS bpFog>GFP | Farley lab | N/A |
| Plasmid: BraS - ZEE bpFog>GFP | This paper | N/A |
| Plasmid: BraS Rand bpFog>GFP | This paper | N/A |
| Plasmid: BraS - FoxA bpFog>GFP | This paper | N/A |
| Plasmid: BraS - Bra bpFog>GFP | This paper | N/A |
| Plasmid: Lama1 bpFog>GFP | This paper | N/A |
| Plasmid: Lama1 bpFog>GFP | This paper | N/A |
| Plasmid: Lama1 - E3 bpFog>GFP | This paper | N/A |
| Plasmid: Lama1 - Z bpFog>GFP | This paper | N/A |
| Plasmid: Lama1 RE3 bpFog>GFP | This paper | N/A |
| Software and Algorithms |  |  |
| Python (version 3.8.6) | Python Software Foundation | https://www.python.org |
| Conda (version 4.9.2) | Anaconda, Inc. | https://docs.conda.io/projects/conda/en/latest/ |
| Bioconda | Grüning et al., 2018 | https://bioconda.github.io |
| Biopython (version 1.78) | Cock et al., 2009 | https://biopython.org |
| FastQC (version 0.11.9) | Babraham Bioinformatics, Babraham Institute | https://www.bioinformatics.babraham.ac.uk/projects/fastqc/ |
| MultiQC (version 1.8) | Ewels et al., 2016 | https://multiqc.info |
| FLASH (version 1.2.11) | Magoč et al., 2011 | http://www.cbcb.umd.edu/software/flash |
| pandas (version 1.2.1) | NumFOCUS | https://pandas.pydata.org |
| NumPy (version 1.20.3) | Harris et al., 2020 | https://numpy.org |
| Matplotlib (version 3.2.2) | Hunter, 2007 | https://matplotlib.org/stable/index.html |
| scikit-learn (version 0.24.1) | Pedregosa et al., 2011 | https://scikit-learn.org/stable/index.html |
| seaborn (version 0.11.1) | Waskom et al., 2021 | https://seaborn.pydata.org/index.html |
| Diverse-Logics-Notochord-Study | Code used in this paper | https://github.com/mragsac/Diverse-Logics-Notochord-Study |

## RESOURCE AVAILABILITY

### Lead contact

Further information and requests for resources and reagents should be directed to and will be fulfilled by the lead contact, Emma Farley.

### Materials availability

Plasmids generated in this study will be deposited in Addgene by the date of publication.

### Data and code availability

- Microscopy and scoring data reported in this paper will be shared by the lead contact upon request.
- All ZEE screen sequencing data will be deposited to GEO and will be made publicly available as of the date of publication. DOIs will be listed in the key resources table.
- All original code will be deposited to GitHub (https://github.com/mragsac/Diverse-Logics-Notochord-Study) and will be made publicly available as of the date of publication. DOIs will be listed in the key resources table.
- Any additional information required to reanalyze the data reported in this paper is available from the lead contact upon request.

## EXPERIMENTAL MODEL AND SUBJECT DETAILS

### Tunicates

Adult *C. intestinalis* type A aka *Ciona robusta* (obtained from M-Rep) were maintained under constant illumination in seawater (obtained from Reliant Aquariums) at 18C. *Ciona* are hermaphroditic, therefore there is only one possible sex for individuals. Age or developmental stage of the embryos studied are indicated in the main text.

## Method Details

### Library Construction

The genomic regions were ordered from Agilent Technologies with adapters containing BseRI sites. This was cloned into the custom-designed SEL-Seq (Synthetic Enhancer Library-Sequencing) vector using type II restriction enzyme BseRI. After cloning, the library was transformed into bacteria (MegaX DHB10 electrocompetent cells), and the culture was grown up until an OD of 1 was reached. DNA was extracted using the Macherey-Nagel Nucleobond Xtra Midi kit. A 30bp barcode with adapters containing Esp3I sites was cloned into this library using type II restriction enzyme Esp3I. The library was transformed into bacteria (MegaX DHB10 electrocompetent cells) and grown up until an OD of 2 was reached. The DNA library was extracted from the bacteria using the Macherey-Nagel Nucleobond Xtra Midi kit.

### Electroporation

Adult *C. intestinalis* type A also known as *Ciona robusta* were obtained from M-Rep and maintained under constant illumination in seawater (obtained from Reliant Aquariums) at 18°C. Dechorionation, *in vitro* fertilization, and electroporation were performed as described previously in Farley et al., 2016. 70 μg DNA was resuspended in 100 μL water and added to 400 μL of 0.96 M D-mannitol. Typically for each electroporation, eggs and sperm were collected from 10 adults. Embryos were fixed at the appropriate developmental stage for 15 minutes in 3.7% formaldehyde. The tissue was then cleared in a series of washes of 0.3% Triton-X in PBS and then of 0.01% Triton-X in PBS. Samples were mounted in Prolong Gold. GFP images were obtained with an Olympus FV3000, using the 40X objective. All constructs were electroporated in three biological replicates.

### ZEE screen

50 μg of the ZEE library was electroporated into ∼5000 fertilized eggs. Embryos developed until 5hrs 30 min at 22°C. Embryos put into TriZol, and RNA was extracted following the manufacturer’s instructions (Life Technologies). The RNA was DNase treated using Turbo DnaseI from Ambion following standard instructions. Poly-A selection was used to obtain only mRNA using poly-A biotinylated beads as per instructions (Dynabeads Life technologies). The mRNA was used in an RT reaction that was specifically selected for the barcoded mRNA (Transcriptor High Fidelity Roche). The RT product was PCR amplified and size selected using Agencourt AMPure beads (Beckman Coulter), then checked for quality and size on the 2100 Bioanalyzer (Agilent) and sent for sequencing on the NovaSeq S4 PE100 mode (Illumina). Three biological replicates were sent for sequencing.

The DNA was extracted by mixing the phenol-chloroform and interphase of TriZol extraction with 500uL of Back Extraction Buffer (4M guanidine thiocyanate, 50mM sodium citrate, and 1M Tris-base). DNA was treated with RnaseA (Thermo Fisher). DNA was cleaned up with phenol:chloroform:isoamyl alcohol (25:24:1) (Life Technologies). The DNA was PCR amplified and size selected using Agencourt AMPure beads (Beckman Coulter), then checked for quality and size on the 2100 Bioanalyzer (Agilent) and sent for sequencing on the NovaSeq S4 PE100 mode (Illumina). Three biological replicates were sent for sequencing.

### Counting Embryos

For each experiment, once embryos had been mounted on slides, slide labels were covered with thick tape and randomly numbered by a laboratory member not involved in this project. Expression of GFP within embryos on each slides was counted blind. In each experiment, all comparative constructs were present, along with a slide with BraS as a reference. Fifty embryos were counted for each biological replicate.

### Acquisition of Images

For enhancers being compared, images were taken from electroporations performed on the same day using identical settings. For representative images, embryos were chosen that represented the average from counting data. All images are subsequently cropped to an appropriate size. In each figure, the same exposure time for each image is shown to allow direct comparison.

### Identification of Putative Notochord Enhancers

We developed a script that allows for the input of any organism’s genome in the fasta file format. The script first looks for an exact match of one of seven canonical Zic family binding sites and their reverse complements. We used the following sites in our search: CAGCTGTG (Zic1/2/3), CCGCAGT (Zic7/3/1), CCGCAGTC (Zic6), CCCGCTGTG (Zic1), CCAGCTGTG (Zic3), CCGCTGTG (Zic2/ZicC), and CCCGCAGTC (Zic5) as these have been identified as functional in previous studies (Matsumoto et al., 2007; Yagi et al., 2004). Next, we drew a window of 30 bp from either end of the canonical Zic family binding site and determine if there are at least two Ets binding site cores (i.e., either GGAA or GGAT and their respective reverse complement sequences) present within the window. If there are at least two Ets binding site cores present, we gather the full Ets binding site by gathering the 2 bp upstream and downstream of the core and finally calculate the relative binding affinity using PBM data. The location of all regions containing at least a single Zic family binding site and two Ets binding sites are saved as part of the genome search.

### Scoring Relative Affinities of Binding Sites

We calculated the relative ETS binding affinity using the median signal intensity of the universal protein binding microarray (PBM) data for mouse Ets-1 proteins from the UniProbe database (http://thebrain.bwh.harvard.edu/uniprobe/index.php) (Hume et al., 2015). Previous studies have shown that the specificity of ETS family members is highly conserved even from flies to humans (Nitta et al., 2015; Wei et al., 2010), and thus ETS-1 is a good proxy for binding affinity in *Ciona* ETS-1 which has a conserved DNA binding domain. The relative affinity score represents the fold change of median signal intensities of the native 8-mer motifs compared to the optimal 8-mer motifs for optimal Ets, which we defined as the CCGGAAGT motif and its corresponding reverse complement.

### Enhancer to Barcode Assignment & Dictionary Analysis

We constructed a dictionary of unique barcode tag-enhancer pairs by not allowing for any mismatches in the ∼68 bp enhancers in our library and by not allowing barcode tag-enhancer pairs to have a read count of fewer than 150 reads. Additionally, we required all barcode tags to be 29 bp or 30 bp in length. If more than one barcode tag was associated with a single enhancer, we included all associated barcode tags that met the aforementioned barcode length and read count requirements. Within our dictionary, we did not find barcode tags that were matched to multiple enhancers. In total, the dictionary contains 90 enhancers that were uniquely mapped to one or more barcode tags, as well as a total of 640 barcode tag-enhancer pairs. On average, enhancers were associated with ∼6 barcodes.

### SEL-Seq Data Analysis

For the whole embryo library, we sequenced barcode tags from the DNA and RNA libraries on the Illumina HiSeq 4000. Reads that perfectly matched barcode tags in our barcode tag-enhancer dictionary were included in the subsequent analysis. 25,361,744 total reads were generated across three replicates of the DNA library, and 393,873,058 total reads were generated across three replicates of the RNA library.

We extracted all of the read sequences from the sequencing libraries and collapse them based on unique sequences, tabulating the number of times a unique sequence appears in the library. Next, we perform preliminary filtering on the unique sequences, filtering out sequences that (i) have N’s present, (ii) are missing the GFP sequence after our expected location of the barcode tag, (iii) contain a barcode not present as an exact match to our enhancer-barcode tag dictionary, (iv) did not meet the minimum read cutoff of 25 reads. Due to the low complexity of the library, we retained all enhancers, even if they only have a single barcode associated with them. For the preliminary filtering step, all DNA and RNA libraries were processed separately.

We first filter our data to only include the set of barcode tags and enhancers that appear in DNA across all replicates. We then normalize our data into RPM and consolidate the expression for each enhancer by taking the average RPM value across barcode tags. Finally, we calculated the log2(RNA/DNA) value for each enhancer to represent the enhancer’s expression. For determining if an enhancer was active, we calculated an “enhancer activity score.” This score is calculated by averaging the log2(RNA/DNA) value across a given enhancer’s biological replicates.

### Data Visualization

Data analysis and visualizations were created using the Python programming language (version 3.8.6). The majority of visualizations used the seaborn (version 0.11.1) and matplotlib (version 3.2.2) packages. Numerical calculations were conducted with the numpy (version 1.20.3) and scikit-learn (version 0.24.1) packages.

## Supplementary figures

**Figure S1.**
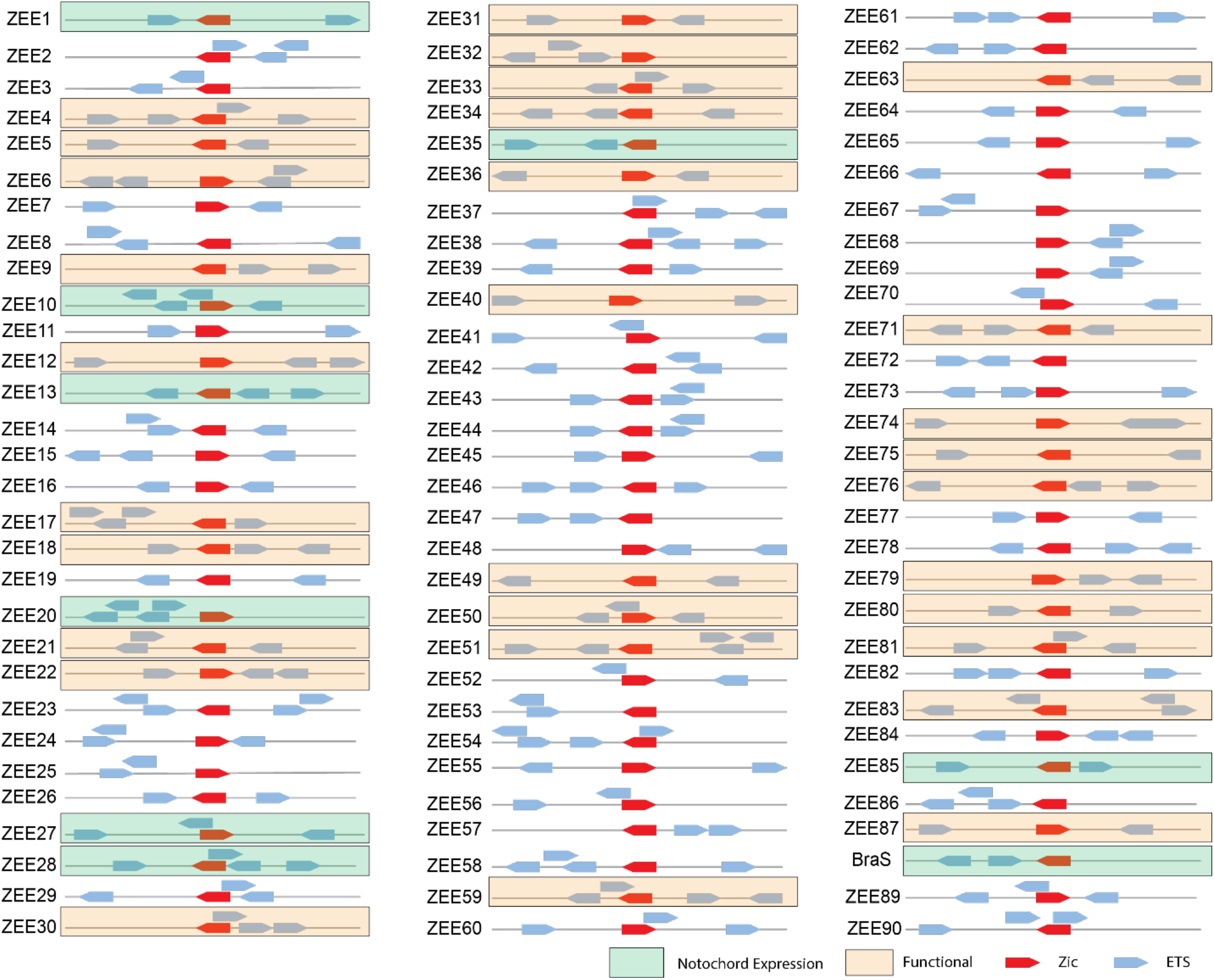
ZEE elements screened. Schematic of each ZEE element. Zic sites are colored red and ETS sites are colored blue. ZEE elements that were functional are boxed in orange. ZEE elements that drove notochord expression are boxed in green.

**Figure S2.**
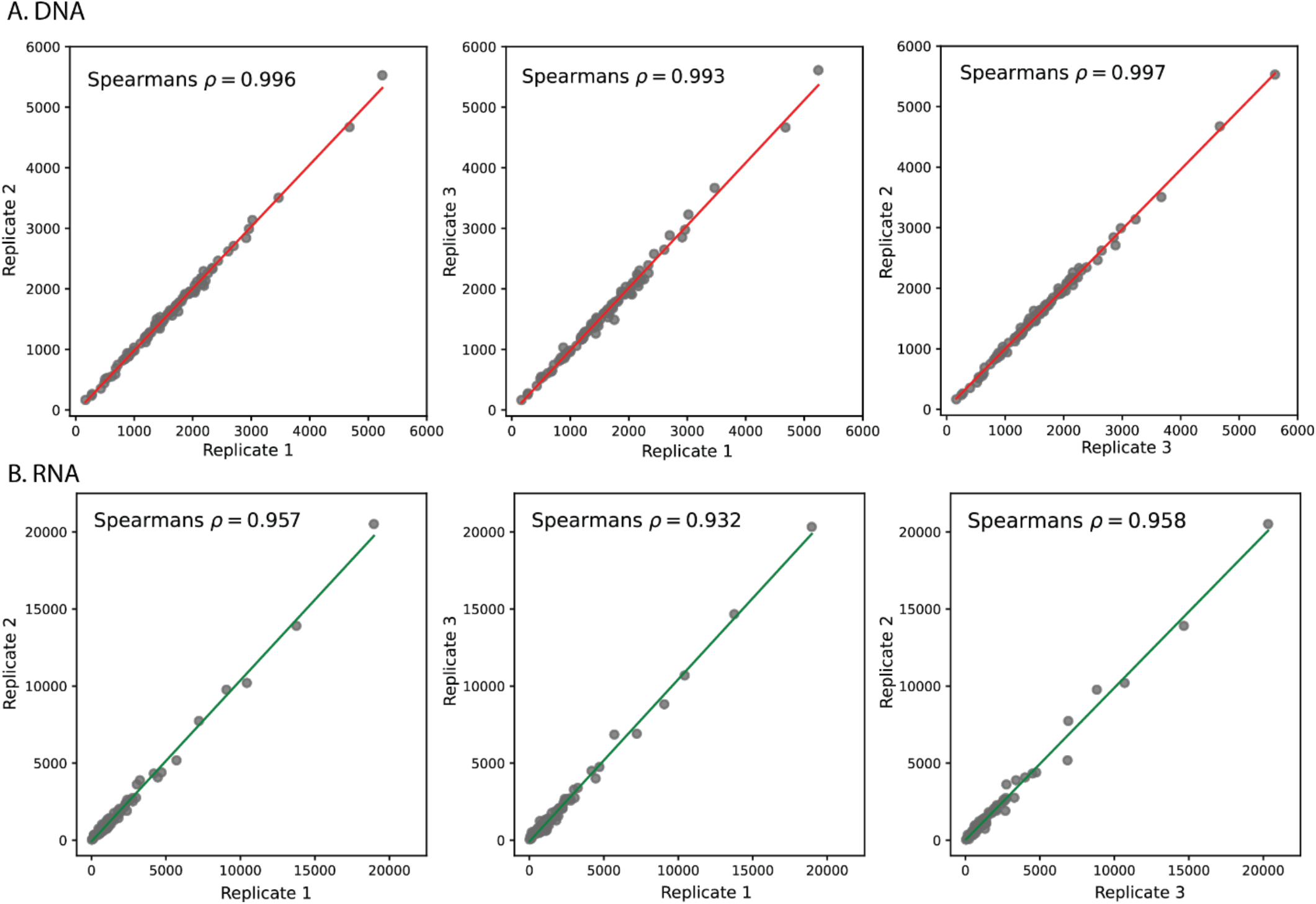
Data quality metrics suggest high robustness of ZEE genomic screen. A. Correlation of DNA plasmids detected between replicates was plotted. All Spearman correlations between replicates were >0.99. B. Correlation of mRNA barcodes detected between replicates was plotted. All Spearman correlations between replicates were >0.9.

**Figure S3.**
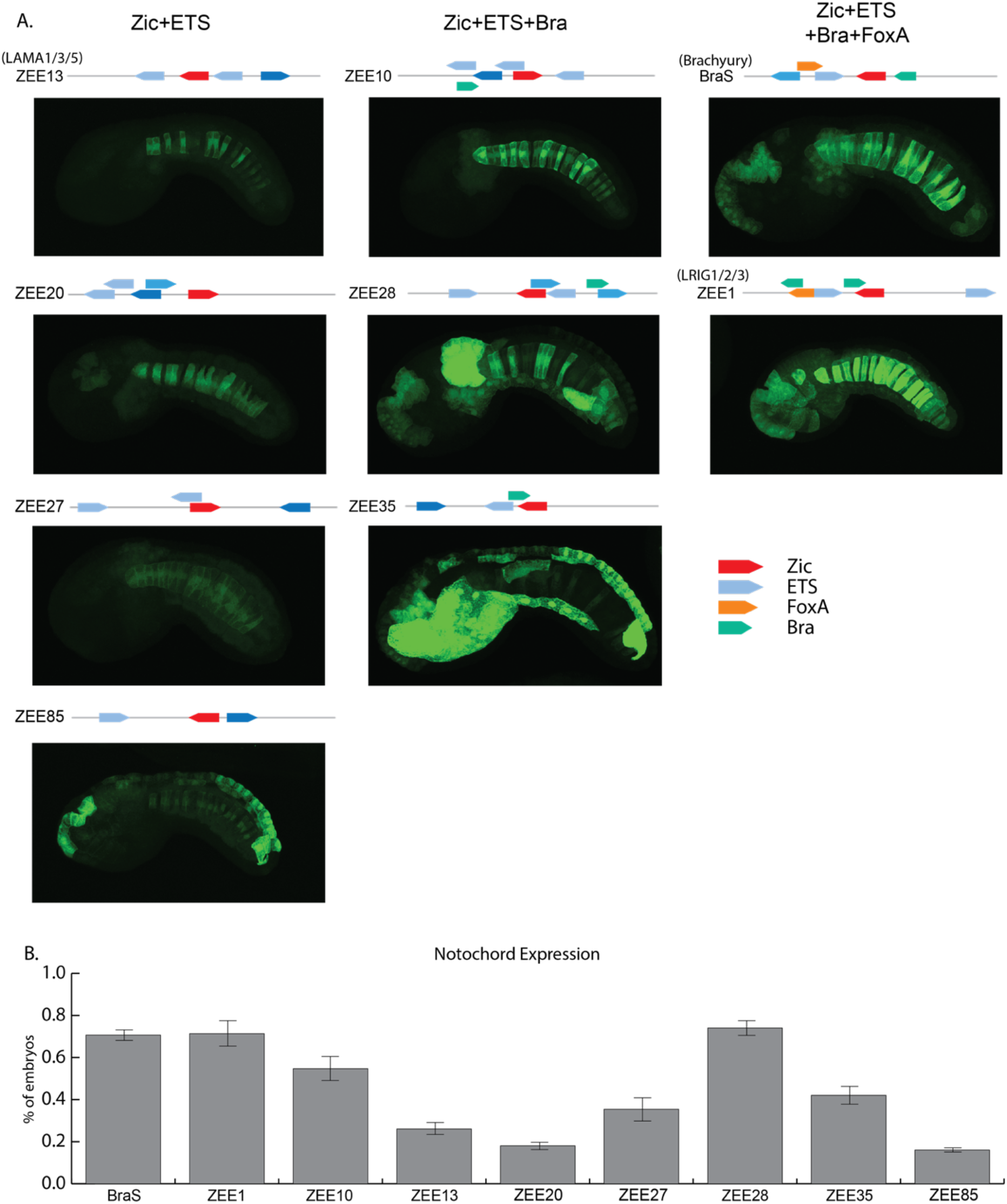
Nine ZEE elements drive notochord expression. A. Images and schematics of the nine notochord enhancers in the ZEE library. Zic (red), ETS (blue), FoxA (orange), and Bra sites (green) are annotated. B. Scoring of notochord expression for embryos electroporated with notochord enhancers. Color saturation of ETS sites indicates binding site affinity (increasing affinity is indicated by an increase in color saturation).

**Figure S4.**
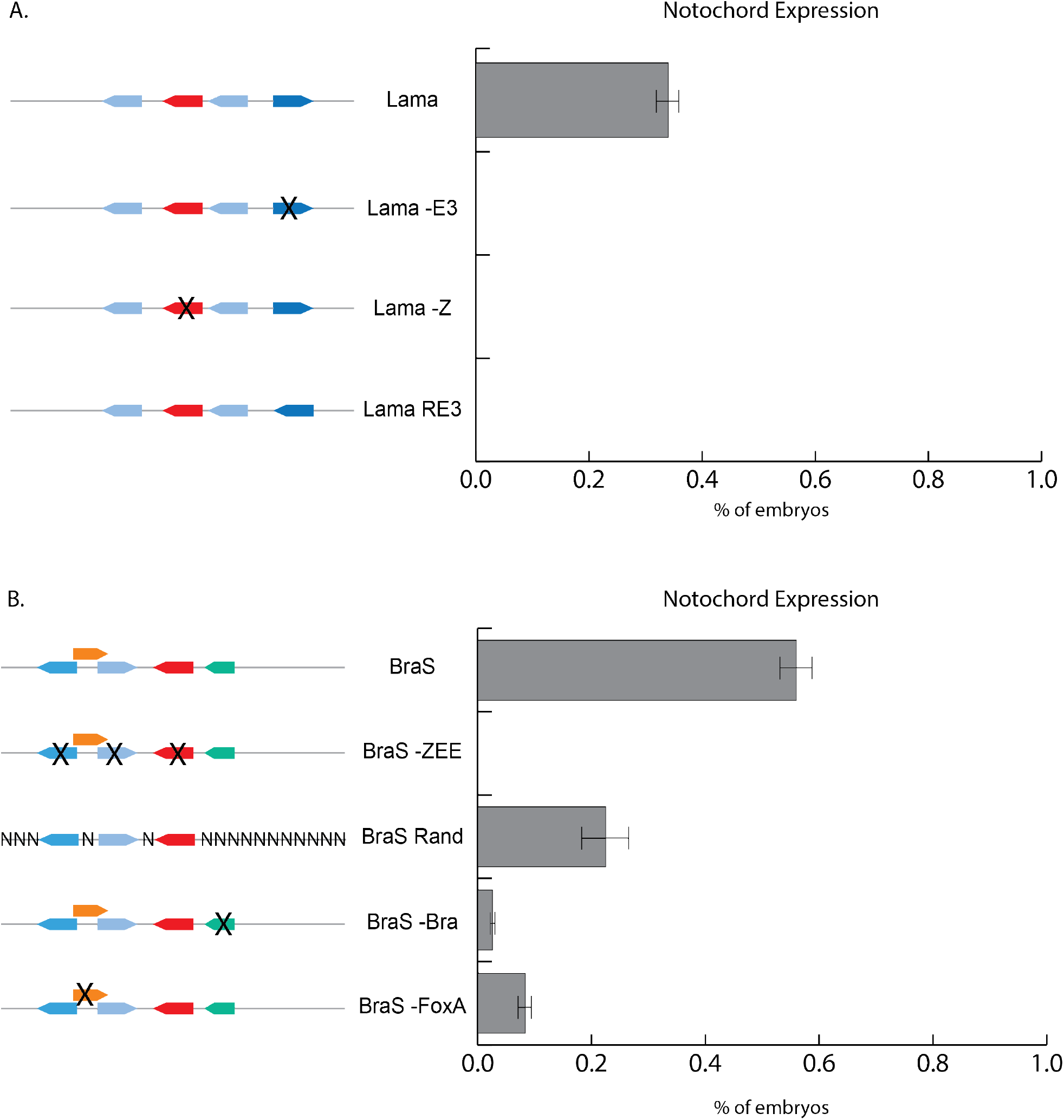
Scoring of manipulated notochord enhancers. A. Scoring of notochord expression for embryos electroporated with the *laminin alpha* (Lama) enhancer, Lama - E3, Lama -Z, and Lama RE3. Lama -E3, Lama -Z, and Lama RE3 all show no notochord expression. B. Scoring of notochord expression for embryos electroporated with Brachyury Shadow (BraS), BraS -ZEE, BraS Rand, BraS -Bra, and BraS – FoxA. BraS - ZEE, BraS Rand, BraS -Bra, and BraS – FoxA all show reduced to no notochord expression compared to BraS. Color saturation of ETS sites indicates binding site affinity (increasing affinity is indicated by an increase in color saturation).

**Figure S5.**
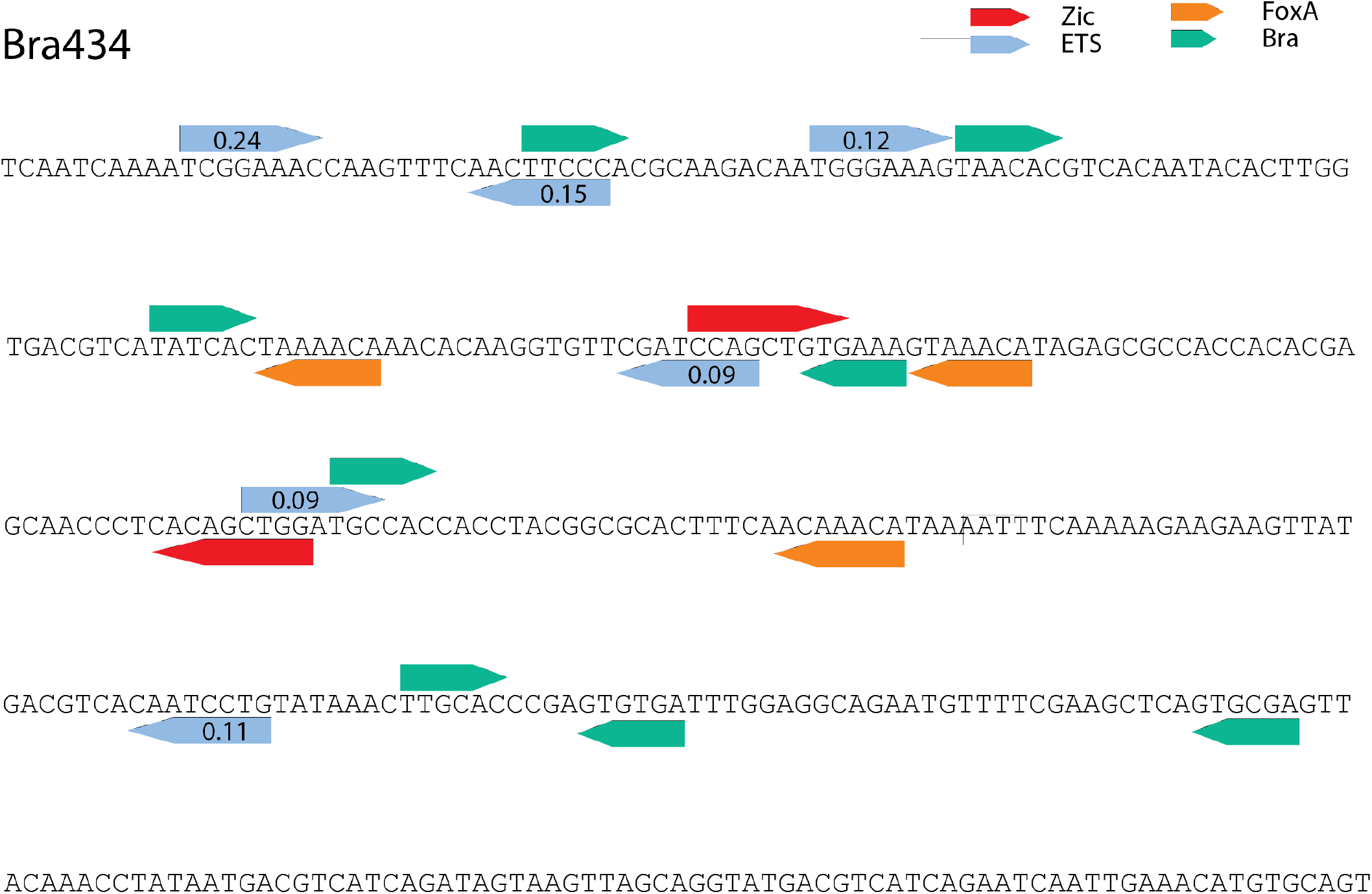
Updated annotation of Bra434. Using PBM, EMSA, and crystal structure data, we propose a new annotation of the Bra434 enhancer. Zic sites in red, ETS sites in light blue, FoxA sites in orange, and Bra sites in green. Affinities of ETS calculated from PBM data (Wei et al., 2010) are labeled.

**Supplementary Table S1: All ZEE elements screened.** This table provides information about all ZEE elements: whether they were tested individually, their enhancer activity score, their genomic location, and their sequence.

**Supplementary Table S2: Scoring of ZEE elements individually tested.** This table provides scoring data for all three replicates of all ZEE elements chosen to be screened individually. Embryos were scored for a6.5, b6.5, notochord, mesenchyme, and endoderm expression.

**Supplementary Table S3: Scoring of manipulations on Lama and BraS enhancers.** This table provides scoring data for the manipulations of the Lama and BraS enhancers. Embryos were scored for notochord expression.

**Supplementary Table S4: Oligonucleotides for Lama and BraS manipulations.** This table provides sequences for oligonucleotides used to mutagenize the Lama and BraS enhancers.

**Supplementary Table S5: Vertebrate enhancers referenced in this study.** This table provides genomic locations of vertebrate enhancers referenced in this study.

## References

Ang, S.-L., Rossant, J., 1994. HNF-3β is essential for node and notochord formation in mouse development. Cell 78, 561–574. https://doi.org/10.1016/0092-8674(94)90522-3

Antosova, B., Smolikova, J., Klimova, L., Lachova, J., Bendova, M., Kozmikova, I., Machon, O., Kozmik, Z., 2016. The Gene Regulatory Network of Lens Induction Is Wired through Meis-Dependent Shadow Enhancers of Pax6. PLoS Genet. 12, e1006441. https://doi.org/10.1371/journal.pgen.1006441

Arnone, M.I., Davidson, E.H., 1997. The hardwiring of development: organization and function of genomic regulatory systems. Dev. Camb. Engl. 124, 1851–1864. https://doi.org/10.1242/dev.124.10.1851

Barolo, S., 2016. How to tune an enhancer. Proc. Natl. Acad. Sci. 113, 6330–6331. https://doi.org/10.1073/pnas.1606109113

Casey, E.S., O’Reilly, M.A., Conlon, F.L., Smith, J.C., 1998. The T-box transcription factor Brachyury regulates expression of eFGF through binding to a non-palindromic response element. Development 125, 3887–3894. https://doi.org/10.1242/dev.125.19.3887

Chesley, P., 1935. Development of the short-tailed mutant in the house mouse. J. Exp. Zool. 70, 429–459. https://doi.org/10.1002/jez.1400700306

Chiba, S., Jiang, D., Satoh, N., Smith, W.C., 2009. brachyury null mutant-induced defects in juvenile ascidian endodermal organs. Development 136, 35–39. https://doi.org/10.1242/dev.030981

Conlon, F.L., Fairclough, L., Price, B.M.J., Casey, E.S., Smith, J.C., 2001. Determinants of T box protein specificity. Development 128, 3749–3758. https://doi.org/10.1242/dev.128.19.3749

Corbo, J.C., Levine, M., Zeller, R.W., 1997. Characterization of a notochord-specific enhancer from the Brachyury promoter region of the ascidian, Ciona intestinalis. Dev. Camb. Engl. 124, 589–602. https://doi.org/10.1242/dev.124.3.589

Dal-Pra, S., Thisse, C., Thisse, B., 2011. FoxA transcription factors are essential for the development of dorsal axial structures. Dev. Biol. 350, 484–495. https://doi.org/10.1016/j.ydbio.2010.12.018

Davidson, B., Christiaen, L., 2006. Linking Chordate Gene Networks to Cellular Behavior in Ascidians. Cell 124, 247–250. https://doi.org/10.1016/j.cell.2006.01.013

Delsuc, F., Brinkmann, H., Chourrout, D., Philippe, H., 2006. Tunicates and not cephalochordates are the closest living relatives of vertebrates. Nature 439, 965–968. https://doi.org/10.1038/nature04336

Di Gregorio, A., Levine, M., 1999. Regulation of Ci-tropomyosin-like, a Brachyury target gene in the ascidian, Ciona intestinalis. Dev. Camb. Engl. 126, 5599–5609. https://doi.org/10.1242/dev.126.24.5599

Dunn, M.P., Di Gregorio, A., 2009. The evolutionarily conserved leprecan gene: Its regulation by Brachyury and its role in the developing Ciona notochord. Dev. Biol. 328, 561–574. https://doi.org/10.1016/j.ydbio.2009.02.007

Elms, P., Scurry, A., Davies, J., Willoughby, C., Hacker, T., Bogani, D., Arkell, R., 2004. Overlapping and distinct expression domains of Zic2 and Zic3 during mouse gastrulation. Gene Expr. Patterns 4, 505–511. https://doi.org/10.1016/j.modgep.2004.03.003

Farley, E.K., Olson, K.M., Zhang, W., Rokhsar, D.S., Levine, M.S., 2016. Syntax compensates for poor binding sites to encode tissue specificity of developmental enhancers. Proc. Natl. Acad. Sci. 113, 6508–6513. https://doi.org/10.1073/pnas.1605085113

Frankel, N., Davis, G.K., Vargas, D., Wang, S., Payre, F., Stern, D.L., 2010. Phenotypic robustness conferred by apparently redundant transcriptional enhancers. Nature 466, 490–493. https://doi.org/10.1038/nature09158

Fujiwara, S., Corbo, J.C., Levine, M., 1998. The snail repressor establishes a muscle/notochord boundary in the Ciona embryo. Development 125, 2511–2520. https://doi.org/10.1242/dev.125.13.2511

Harvey, S.A., Tümpel, S., Dubrulle, J., Schier, A.F., Smith, J.C., 2010. no tail integrates two modes of mesoderm induction. Development 137, 1127–1135. https://doi.org/10.1242/dev.046318

Heinz, S., Benner, C., Spann, N., Bertolino, E., Lin, Y.C., Laslo, P., Cheng, J.X., Murre, C., Singh, H., Glass, C.K., 2010. Simple combinations of lineage-determining transcription factors prime cis-regulatory elements required for macrophage and B cell identities. Mol. Cell 38, 576–589. https://doi.org/10.1016/j.molcel.2010.05.004

Herrmann, B.G., Kispert, A., 1994. The T genes in embryogenesis. Trends Genet. TIG 10, 280–286. https://doi.org/10.1016/0168-9525(90)90011-t

Hong, J.-W., Hendrix, D.A., Levine, M.S., 2008. Shadow enhancers as a source of evolutionary novelty. Science 321, 1314. https://doi.org/10.1126/science.1160631

Hudson, C., Lotito, S., Yasuo, H., 2007. Sequential and combinatorial inputs from Nodal, Delta2/Notch and FGF/MEK/ERK signalling pathways establish a grid-like organisation of distinct cell identities in the ascidian neural plate. Dev. Camb. Engl. 134, 3527–3537. https://doi.org/10.1242/dev.002352

Hudson, C., Sirour, C., Yasuo, H., 2016. Co-expression of Foxa.a, Foxd and Fgf9/16/20 defines a transient mesendoderm regulatory state in ascidian embryos. eLife 5, e14692. https://doi.org/10.7554/eLife.14692

Hume, M.A., Barrera, L.A., Gisselbrecht, S.S., Bulyk, M.L., 2015. UniPROBE, update 2015: new tools and content for the online database of protein-binding microarray data on protein–DNA interactions. Nucleic Acids Res. 43, D117–D122. https://doi.org/10.1093/nar/gku1045

Ikeda, T., Satou, Y., 2016. Differential temporal control of Foxa.a and Zic-r.b specifies brain versus notochord fate in the ascidian embryo. Development dev.142174. https://doi.org/10.1242/dev.142174

Imai, K.S., Hino, K., Yagi, K., Satoh, N., Satou, Y., 2004. Gene expression profiles of transcription factors and signaling molecules in the ascidian embryo: towards a comprehensive understanding of gene networks. Development 131, 4047–4058. https://doi.org/10.1242/dev.01270

Imai, K.S., Levine, M., Satoh, N., Satou, Y., 2006. Regulatory Blueprint for a Chordate Embryo. Science 312, 1183–1187. https://doi.org/10.1126/science.1123404

Imai, K.S., Satoh, N., Satou, Y., 2002a. Early embryonic expression of FGF4/6/9 gene and its role in the induction of mesenchyme and notochord in Ciona savignyi embryos. Development 129, 1729–1738. https://doi.org/10.1242/dev.129.7.1729

Imai, K.S., Satou, Y., Satoh, N., 2002b. Multiple functions of a Zic-like gene in the differentiation of notochord, central nervous system and muscle in Ciona savignyi embryos. Dev. Camb. Engl. 129, 2723–2732. https://doi.org/10.1242/dev.129.11.2723

Jiang, D., Smith, W.C., 2007. Ascidian notochord morphogenesis. Dev. Dyn. Off. Publ. Am. Assoc. Anat. 236, 1748–1757. https://doi.org/10.1002/dvdy.21184

José-Edwards, D.S., Oda-Ishii, I., Kugler, J.E., Passamaneck, Y.J., Katikala, L., Nibu, Y., Di Gregorio, A., 2015. Brachyury, Foxa2 and the cis-Regulatory Origins of the Notochord. PLoS Genet. 11, e1005730. https://doi.org/10.1371/journal.pgen.1005730

Katikala, L., Aihara, H., Passamaneck, Y.J., Gazdoiu, S., José-Edwards, D.S., Kugler, J.E., Oda-Ishii, I., Imai, J.H., Nibu, Y., Di Gregorio, A., 2013. Functional Brachyury binding sites establish a temporal read-out of gene expression in the Ciona notochord. PLoS Biol. 11, e1001697. https://doi.org/10.1371/journal.pbio.1001697

Kumano, G., Yamaguchi, S., Nishida, H., 2006. Overlapping expression of FoxA and Zic confers responsiveness to FGF signaling to specify notochord in ascidian embryos. Dev. Biol. 300, 770–784. https://doi.org/10.1016/j.ydbio.2006.07.033

Levine, M., 2010. Transcriptional Enhancers in Animal Development and Evolution. Curr. Biol. 20, R754–R763. https://doi.org/10.1016/j.cub.2010.06.070

Levo, M., Segal, E., 2014. In pursuit of design principles of regulatory sequences. Nat. Rev. Genet. 15, 453–468. https://doi.org/10.1038/nrg3684

Li, J., Dantas Machado, A.C., Guo, M., Sagendorf, J.M., Zhou, Z., Jiang, L., Chen, X., Wu, D., Qu, L., Chen, Z., Chen, L., Rohs, R., Chen, Y., 2017. Structure of the Forkhead Domain of FOXA2 Bound to a Complete DNA Consensus Site. Biochemistry 56, 3745–3753. https://doi.org/10.1021/acs.biochem.7b00211

Liu, F., Posakony, J.W., 2012. Role of Architecture in the Function and Specificity of Two Notch-Regulated Transcriptional Enhancer Modules. PLoS Genet. 8, e1002796. https://doi.org/10.1371/journal.pgen.1002796

Lolas, M., Valenzuela, P.D.T., Tjian, R., Liu, Z., 2014. Charting Brachyury-mediated developmental pathways during early mouse embryogenesis. Proc. Natl. Acad. Sci. 111, 4478–4483. https://doi.org/10.1073/pnas.1402612111

Machingo, Q.J., Fritz, A., Shur, B.D., 2006. A beta1,4-galactosyltransferase is required for convergent extension movements in zebrafish. Dev. Biol. 297, 471–482. https://doi.org/10.1016/j.ydbio.2006.05.024

Matsumoto, J., Kumano, G., Nishida, H., 2007. Direct activation by Ets and Zic is required for initial expression of the Brachyury gene in the ascidian notochord. Dev. Biol. 306, 870–882. https://doi.org/10.1016/j.ydbio.2007.03.034

Maurano, M.T., Humbert, R., Rynes, E., Thurman, R.E., Haugen, E., Wang, H., Reynolds, A.P., Sandstrom, R., Qu, H., Brody, J., Shafer, A., Neri, F., Lee, K., Kutyavin, T., Stehling-Sun, S., Johnson, A.K., Canfield, T.K., Giste, E., Diegel, M., Bates, D., Hansen, R.S., Neph, S., Sabo, P.J., Heimfeld, S., Raubitschek, A., Ziegler, S., Cotsapas, C., Sotoodehnia, N., Glass, I., Sunyaev, S.R., Kaul, R., Stamatoyannopoulos, J.A., 2012. Systematic localization of common disease-associated variation in regulatory DNA. Science 337, 1190–1195. https://doi.org/10.1126/science.1222794

Miya, T., Nishida, H., 2003. An Ets transcription factor, HrEts, is target of FGF signaling and involved in induction of notochord, mesenchyme, and brain in ascidian embryos. Dev. Biol. 261, 25–38. https://doi.org/10.1016/s0012-1606(03)00246-x

Müller, C.W., Herrmann, B.G., 1997. Crystallographic structure of the T domain–DNA complex of the Brachyury transcription factor. Nature 389, 884–888. https://doi.org/10.1038/39929

Nitta, K.R., Jolma, A., Yin, Y., Morgunova, E., Kivioja, T., Akhtar, J., Hens, K., Toivonen, J., Deplancke, B., Furlong, E.E.M., Taipale, J., 2015. Conservation of transcription factor binding specificities across 600 million years of bilateria evolution. eLife 4, e04837. https://doi.org/10.7554/eLife.04837

Osterwalder, M., Barozzi, I., Tissières, V., Fukuda-Yuzawa, Y., Mannion, B.J., Afzal, S.Y., Lee, E.A., Zhu, Y., Plajzer-Frick, I., Pickle, C.S., Kato, M., Garvin, T.H., Pham, Q.T., Harrington, A.N., Akiyama, J.A., Afzal, V., Lopez-Rios, J., Dickel, D.E., Visel, A., Pennacchio, L.A., 2018. Enhancer redundancy provides phenotypic robustness in mammalian development. Nature 554, 239–243. https://doi.org/10.1038/nature25461

Parsons, M.J., Campos, I., Hirst, E.M.A., Stemple, D.L., 2002. Removal of dystroglycan causes severe muscular dystrophy in zebrafish embryos. Dev. Camb. Engl. 129, 3505–3512. https://doi.org/10.1242/dev.129.14.3505

Passamaneck, Y.J., Katikala, L., Perrone, L., Dunn, M.P., Oda-Ishii, I., Di Gregorio, A., 2009. Direct activation of a notochord cis-regulatory module by Brachyury and FoxA in the ascidian Ciona intestinalis. Dev. Camb. Engl. 136, 3679–3689. https://doi.org/10.1242/dev.038141

Perry, M.W., Boettiger, A.N., Bothma, J.P., Levine, M., 2010. Shadow enhancers foster robustness of Drosophila gastrulation. Curr. Biol. CB 20, 1562–1567. https://doi.org/10.1016/j.cub.2010.07.043

Picco, V., Hudson, C., Yasuo, H., 2007. Ephrin-Eph signalling drives the asymmetric division of notochord/neural precursors in Ciona embryos. Dev. Camb. Engl. 134, 1491–1497. https://doi.org/10.1242/dev.003939

Pijuan-Sala, B., Wilson, N.K., Xia, J., Hou, X., Hannah, R.L., Kinston, S., Calero-Nieto, F.J., Poirion, O., Preissl, S., Liu, F., Göttgens, B., 2020. Single-cell chromatin accessibility maps reveal regulatory programs driving early mouse organogenesis. Nat. Cell Biol. 22, 487–497. https://doi.org/10.1038/s41556-020-0489-9

Pollard, S.M., Parsons, M.J., Kamei, M., Kettleborough, R.N.W., Thomas, K.A., Pham, V.N., Bae, M.-K., Scott, A., Weinstein, B.M., Stemple, D.L., 2006. Essential and overlapping roles for laminin alpha chains in notochord and blood vessel formation. Dev. Biol. 289, 64–76. https://doi.org/10.1016/j.ydbio.2005.10.006

Reeves, W.M., Shimai, K., Winkley, K.M., Veeman, M.T., 2021. Brachyury controls Ciona notochord fate as part of a feed-forward network. Dev. Camb. Engl. 148, dev195230. https://doi.org/10.1242/dev.195230

Reeves, W.M., Wu, Y., Harder, M.J., Veeman, M.T., 2017. Functional and evolutionary insights from the Ciona notochord transcriptome. Dev. Camb. Engl. 144, 3375–3387. https://doi.org/10.1242/dev.156174

Schifferl, D., Scholze-Wittler, M., Wittler, L., Veenvliet, J.V., Koch, F., Herrmann, B.G., 2021. A 37 kb region upstream of brachyury comprising a notochord enhancer is essential for notochord and tail development. Development 148, dev200059. https://doi.org/10.1242/dev.200059

Schulte-Merker, S., Smith, J.C., 1995. Mesoderm formation in response to Brachyury requires FGF signalling. Curr. Biol. 5, 62–67. https://doi.org/10.1016/S0960-9822(95)00017-0

Scott, A., Stemple, D.L., 2005. Zebrafish notochordal basement membrane: signaling and structure. Curr. Top. Dev. Biol. 65, 229–253. https://doi.org/10.1016/S0070-2153(04)65009-5

Shimai, K., Veeman, M., 2021. Quantitative Dissection of the Proximal Ciona brachyury Enhancer. Front. Cell Dev. Biol. 9, 804032. https://doi.org/10.3389/fcell.2021.804032

Small, S., Blair, A., Levine, M., 1992. Regulation of even-skipped stripe 2 in the Drosophila embryo. EMBO J. 11, 4047–4057. https://doi.org/10.1002/j.1460-2075.1992.tb05498.x

Spitz, F., Furlong, E.E.M., 2012. Transcription factors: from enhancer binding to developmental control. Nat. Rev. Genet. 13, 613–626. https://doi.org/10.1038/nrg3207

Stemple, D.L., 2005. Structure and function of the notochord: an essential organ for chordate development. Development 132, 2503–2512. https://doi.org/10.1242/dev.01812

Swanson, C.I., Evans, N.C., Barolo, S., 2010. Structural Rules and Complex Regulatory Circuitry Constrain Expression of a Notch- and EGFR-Regulated Eye Enhancer. Dev. Cell 18, 359–370. https://doi.org/10.1016/j.devcel.2009.12.026

Tak, Y.G., Farnham, P.J., 2015. Making sense of GWAS: using epigenomics and genome engineering to understand the functional relevance of SNPs in non-coding regions of the human genome. Epigenetics Chromatin 8, 57. https://doi.org/10.1186/s13072-015-0050-4

Takahashi, H., Mitani, Y., Satoh, G., Satoh, N., 1999. Evolutionary alterations of the minimal promoter for notochord-specific Brachyury expression in ascidian embryos. Dev. Camb. Engl. 126, 3725–3734. https://doi.org/10.1242/dev.126.17.3725

Thanos, D., Maniatis, T., 1995. Virus induction of human IFN beta gene expression requires the assembly of an enhanceosome. Cell 83, 1091–1100. https://doi.org/10.1016/0092-8674(95)90136-1

Veeman, M.T., Nakatani, Y., Hendrickson, C., Ericson, V., Lin, C., Smith, W.C., 2008. Chongmague reveals an essential role for laminin-mediated boundary formation in chordate convergence and extension movements. Dev. Camb. Engl. 135, 33–41. https://doi.org/10.1242/dev.010892

Visel, A., Rubin, E.M., Pennacchio, L.A., 2009. Genomic views of distant-acting enhancers. Nature 461, 199–205. https://doi.org/10.1038/nature08451

Wagner, E., Levine, M., 2012. FGF signaling establishes the anterior border of the Ciona neural tube. Dev. Camb. Engl. 139, 2351–2359. https://doi.org/10.1242/dev.078485

Wei, G.-H., Badis, G., Berger, M.F., Kivioja, T., Palin, K., Enge, M., Bonke, M., Jolma, A., Varjosalo, M., Gehrke, A.R., Yan, J., Talukder, S., Turunen, M., Taipale, M., Stunnenberg, H.G., Ukkonen, E., Hughes, T.R., Bulyk, M.L., Taipale, J., 2010. Genome-wide analysis of ETS-family DNA-binding in vitro and in vivo. EMBO J. 29, 2147–2160. https://doi.org/10.1038/emboj.2010.106

Weinstein, D.C., Ruiz i Altaba, A., Chen, W.S., Hoodless, P., Prezioso, V.R., Jessell, T.M., Darnell, J.E., 1994. The winged-helix transcription factor HNF-3β is required for notochord development in the mouse embryo. Cell 78, 575–588. https://doi.org/10.1016/0092-8674(94)90523-1

Wilkinson, D.G., Bhatt, S., Herrmann, B.G., 1990. Expression pattern of the mouse T gene and its role in mesoderm formation. Nature 343, 657–659. https://doi.org/10.1038/343657a0

Winkley, K.M., Reeves, W.M., Veeman, M.T., 2021. Single-cell analysis of cell fate bifurcation in the chordate Ciona. BMC Biol. 19, 180. https://doi.org/10.1186/s12915-021-01122-0

Yagi, K., Satou, Y., Satoh, N., 2004. A zinc finger transcription factor, ZicL, is a direct activator of Brachyury in the notochord specification of Ciona intestinalis. Development 131, 1279–1288. https://doi.org/10.1242/dev.01011

Yasuo, H., Hudson, C., 2007. FGF8/17/18 functions together with FGF9/16/20 during formation of the notochord in Ciona embryos. Dev. Biol. 302, 92–103. https://doi.org/10.1016/j.ydbio.2006.08.075

Yasuo, H., Satoh, N., 1993. Function of vertebrate T gene. Nature 364, 582–583. https://doi.org/10.1038/364582b0

